# Interventionally-guided representation learning for robust and interpretable AI models in cancer medicine

**DOI:** 10.1101/2025.07.21.662350

**Authors:** Dom Kirkham, Riccardo Masina, Stephen-John Sammut, Sach Mukherjee, Oscar M. Rueda

**Affiliations:** MRC Biostatistics Unit, University of Cambridge, Cambridge, UK; Cancer Research UK Cambridge Institute, University of Cambridge, Cambridge, UK; Breast Cancer Now Toby Robins Research Centre, The Institute of Cancer Research, London, UK; The Royal Marsden Hospital NHS Foundation Trust, London, UK; German Centre for Neurodegenerative Diseases (DZNE) & University of Bonn, Bonn, Germany

**Author notes:** Contributing authors.

**Keywords:** biomedical AI, generative models, cancer biology, machine learning

## Abstract

Machine learning models hold promise in cancer medicine but often lack robustness and interpretability. We introduce a new class of model for high-dimensional molecular data that incorporate interventional auxiliary information to learn latent representations that are informative and interpretable by design. By using causal signals from genetic loss-of-function screens, our approach generates representations that generalize well across data distributions and biological contexts. In cancer cell line datasets, we show that causal guidance enables “zero-shot” transfer to cancer types unseen during training. Moreover, models trained solely on cell line data translate effectively to clinical cohorts, demonstrating strong “bench-to-bedside” generalization without fine-tuning. This strategy highlights a scalable way to leverage tractable laboratory assays for clinical modeling. More broadly, our results establish how integrating causal biological information within generative frameworks enhances data efficiency, interpretability, and robustness, opening avenues for a new generation of scientifically informed AI models in molecular medicine.

## 1 Introduction

AI and machine learning (ML) methods are increasingly prominent in cancer biology and medicine. One common use of AI and ML models is to predict phenotypes based on high-throughput data. However, recent work in core AI and ML as well as in a number of application domains has highlighted the fact that trained AI models can suffer from a dramatic lack of robustness and struggle to generalise to new contexts, tasks and populations, hence significantly limiting practical utility [1–3]. This is because flexible AI models can learn to rely on patterns in training data which are not consistent across contexts, resulting in poor generalisation and performance in practice [4]. Since biologically meaningful models with suitable inductive biases are likely to generalise better [5–9], these issues are also thought to be related to a lack of interpretability. Improving robustness and generalisation is critical in realising the potential of ML models in cancer, since data generated in real-world scenarios may differ substantially from the data used for model training.

Human biology—and cancer biology in particular—is highly heterogeneous in terms of phenotypes and responses to perturbations such as drug treatment, and this heterogeneity has long challenged research in cancer biology and medicine. At the same time, many shared biological mechanisms underpin observed variation. Such mechanisms, while complex and still not fully understood, are also not arbitrary but rather linked (via diverse processes) to underlying sources of genetic and epigenetic variation. This suggests the hypothesis that viewing high-dimensional molecular data through an appropriate lens that leverages shared mechanisms could enhance generalisability. From an ML perspective, this implies that flexible models that are guided towards the discovery of appropriate representations could provide a way to practically study complex, high-dimensional molecular data.

In this paper, motivated by the preceding, we propose a general class of ML models called *Prediction with interpretable Constraints* or PiCo that leverage causal information to guide representation learning. The key idea is to augment standard neural network-based representation learning and generative modelling with additional information from interventional experiments relevant to the specific setting (cancer biology). Causal and/or interventional information obtained from designed experiments is crucial to scientific modelling since it pushes systems to new configurations via specific interventions, which from an ML point of view corresponds to particularly rich and informative signals by which to guide learning.

In our model, this is done by constraining certain latent variables with information from interventional experiments (as detailed below), which, as we show, leads to models that are both interpretable and able to generalise to very different target settings. The particular information we use is from CRISPR-Cas9 screens. CRISPR-Cas9 screening and its variants offer a scalable method to study responses to genetic perturbation in cells. By inducing the loss-of-function of individual genes across the genome, the dependency of cells on the function of these genes can be quantified via relative changes in cell fitness following the perturbations. We leverage CRISPR-Cas9 screens to provide interpretable constraints on learned representations, which we show improves generalisation compared to baselines. We demonstrate applications of our model which leverage this idea of shared mechanisms of response to perturbation, revealing that placing biologically-informed constraints on models can be important in preventing models from over-fitting to training distributions, thus enabling broader generalisation. PiCo achieves these improvements within a flexible, generative framework and constraints can be placed on the model in a data-driven rather than fixed manner. Our model does not rely on mechanisms being shared exactly but rather on the notion that certain kinds of auxiliary information can provide mechanistic insights to usefully inform ML models during training. Since PiCo generates interpretable and information-rich representations, predictive modelling can finally be carried out using only very simple linear models, ensuring simplicity and data efficiency.

We show that PiCo produces robust and generalisable predictions of response to pharmacological perturbation, including in zero-shot settings where a trained model is applied to entirely new data in an “out-of-the-box” fashion without any further training. In cell lines, this allows PiCo to generalise to entirely new cancer types without any further model training or fine-tuning. Turning to data from clinical cohorts, we show that low-dimensional representations trained using only data from cancer cell lines are robust enough to be directly used on patient data without any further adaptation or fine-tuning.

We demonstrate applications of PiCo to a common scenario in drug response prediction on cancer cell lines—using mRNA gene expression—first learning how gene expression relates to response to genetic perturbation using CRISPR screens as auxiliary data to constrain representations, and then using these learned constrained representations in drug response prediction.

PiCo is related to several strands of recent research in ML and cancer biology. The notion that generative models can gain from the incorporation of prior knowledge is widely appreciated, and our work is related technically to efforts in the ML community to develop constrained variants of generative models such as VAEs that leverage appropriate inductive biases [10–12]. Our work is also related to a line of research on out-of-distribution generalisation, transfer learning, meta learning, and domain generalisation, in which variational autoencoders, among other ML models, are designed to cope better with changes to statistical distributions and tasks [13–17]. Our work complements the preceding by showing how, in a cancer biology setting, specific constraints aimed at capturing shared underlying mechanisms of response to perturbation can provide effective and practically applicable guidance for learning from high-dimensional molecular data. An important line of recent work in AI has involved training generalisable foundation models using very large data [18, 19]. This strategy, while important and effective in many settings, is extraordinarily resource-intensive, both with respect to data and compute. In contrast, our approach uses causal information to permit strongly generalisable learning that is at the same time highly data- and compute-efficient.

In cancer medicine, an increasing amount of interest has been shown in utilising prior knowledge to improve models for prediction tasks. In many cases, this has come in the form of integrating curated knowledge from databases, which can be used to provide an inductive bias to models, either through the use of network structures [20, 21] or by constraining model architectures to relate to biological pathways [5–7]. PiCo combines the positives of both of these approaches—utilising an external dataset for pretraining, while capturing the benefit provided through this pretraining in a consistently interpretable way.

Many recent models for drug response prediction using gene expression or multi-omics rely on deep neural networks to effectively process complex, high-dimensional data [22]. Intending to improve interpretability in deep learning models and their generalisation to new domains, recent works in other areas have introduced *biologically informed* ML methods. This typically involves incorporating prior knowledge from Gene Ontology (GO), protein-protein interaction (PPI) networks, or functional association networks (e.g. STRING [23]) to constrain model architecture [5–7]. While the incorporation of prior biological knowledge in this way can be insightful, the interactions documented can lack relevance to the context being studied. This issue has previously been approached in the context of reference mapping for single-cell RNASeq by allowing the combined use of prior knowledge and interactions learned from data in predictive models [8]. However, prior knowledge-informed models can face limitations due to the fact that current knowledge of signalling pathways and protein interactions is incomplete and likely too rigid. In contrast, our approach is flexible and data-adaptive, taking advantage of constraints based on auxiliary data.

Predicting a target based on interpretable concepts is popular in other areas of AI and ML, notably with *concept-based models* which provide interpretability while improving generalisation to new settings by making predictions based on human-interpretable visual concepts [24, 25].

Settings similar to those studied here are increasingly common in cancer biology due to the growing availability of large multi-omics datasets. One such recent example involves predicting cancer treatment response by imputing transcriptomics from histopathology images, showing that these imputed values are more useful for predicting outcomes than naively learned features [26].

To develop the models and test these hypotheses, we leveraged data from 1019 cell lines with genomewide CRISPR-Cas9 loss-of-function screening information from the DepMap 23Q2 dataset [27] and 686 cell lines with extensive pharmacological screening information from the GDSC2 dataset [28], drawing relationships between gene effect and drug response using 499 cell lines present in both screens. We extended the study by utilising the PiCo framework for prediction of clinical outcomes in two breast cancer cohorts, TransNEO [29] and SCAN-B [30], using ARTemis+PBCP [29] and The Cancer Genome Atlas (TCGA) [31] as external validation sets, respectively, for each study.

## 2 Results

### 2.1 Model and experiments overview

For representation learning using high-throughput data, we developed a semi-supervised variational autoencoder (VAE)-based [13] model suitable for utilising CRISPR gene effect data as auxiliary information to constrain representations. We refer to this generative model as *interpretable Constrained VAE* (iCoVAE). We then used representations generated by iCoVAE in drug response prediction on cancer cell lines, within a framework we refer to as *Prediction with interpretable Constraints* (PiCo). Constraints are imposed by requiring that specific dimensions in the latent space are directly linked to specific features derived from the CRISPR screen—namely the effect of knocking out a specific gene on cell fitness (henceforth referred to as *gene effect*); the remaining dimensions in the latent space are unconstrained and able to capture other variation in the data which is unrelated to the features from the auxiliary data. The iCoVAE builds both upon the semi-supervised variational autoencoder framework [13] and the Characteristic Capturing VAE (CCVAE) [10] (see Section 4 for further details). To study the ability of PiCo to generalise outside of its training distribution (“out-of-distribution” or OOD generalisation), we fitted models using only data from cancer cell lines of a subset of cancer types, then used these models to generate representations for *other* cancer types entirely unseen during training (Fig. B2b). This differs from the classical train-test setting because entire cancer types—and not just individual samples—are unseen during training.

### 2.2 iCoVAE produces rich and interpretable representations of cancer cell line transcriptomics

We first trained iCoVAE on data from cancer cell lines in DepMap to better understand the quality of its generated representations of gene expression data. Fig. 2a demonstrates the richness of these representations, using three different iCoVAE models with constraints from the auxiliary CRISPR screening data selected from a data-derived set of genes related to drug response in the drugs ceralasertib (AZD6738), lapatinib and palbociclib (See Methods). Where metrics are provided, these correspond to a mean across 10 model initialisations, each with different random seeds, unless otherwise specified.

**Fig. 1.**
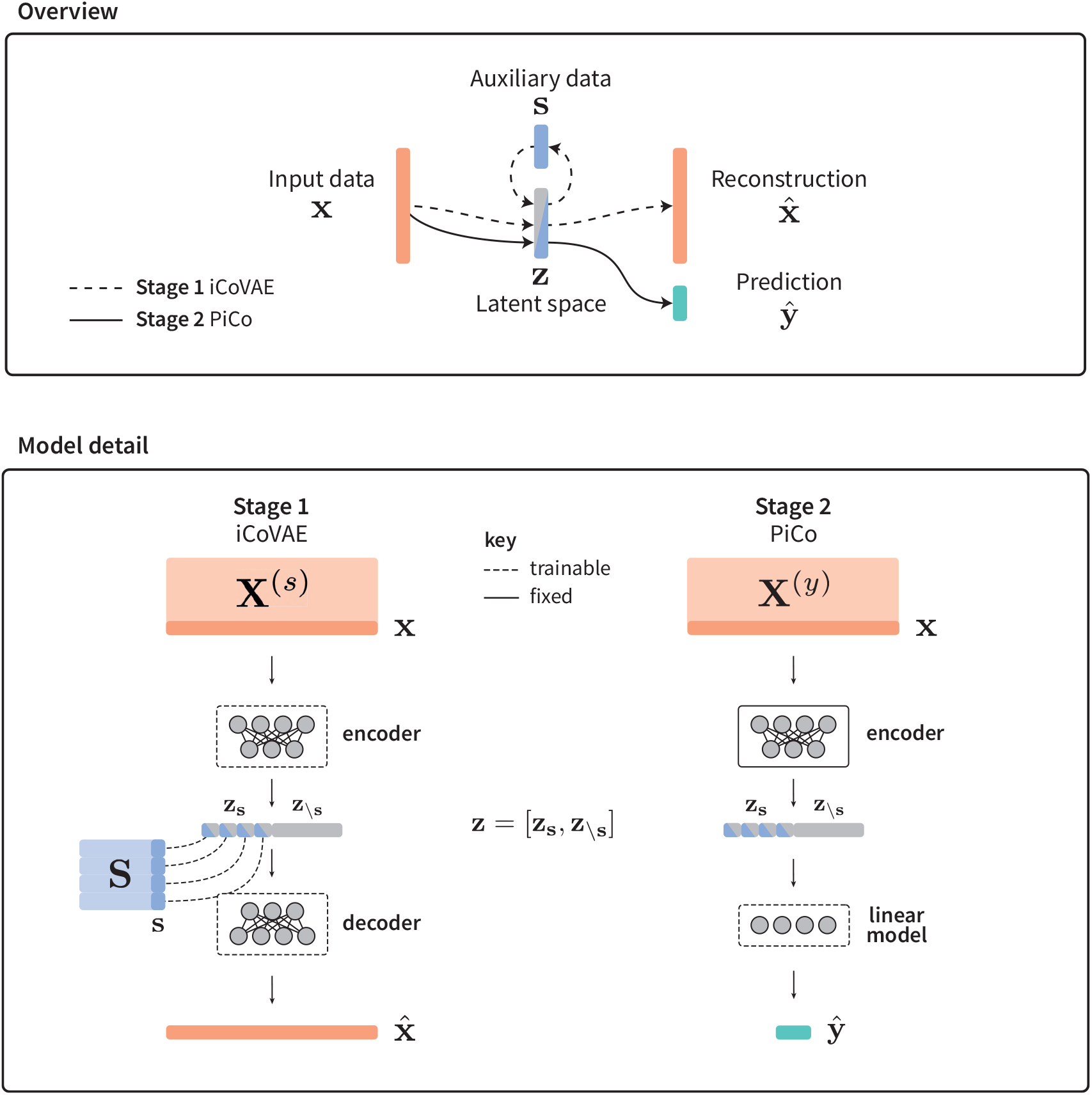
PiCo model. Model overview In Stage 1, A semi-supervised variational autoencoder-based model (iCoVAE) is pretrained using input data **X**^(*s*)^ for which some auxiliary data **S** is available. Features within the constrained latent space **z**_*s*_ are individually constrained using selected features from **S**. In Stage 2, the encoder of the pretrained model is then fixed and used to extract representations for input data **X**^(*y*)^ for which some target **y** is available. These representations are then used to train a linear model to make predictions for the target, **ŷ**. Auxiliary data **s** is not required in Stage 2.

**Fig. 2.**
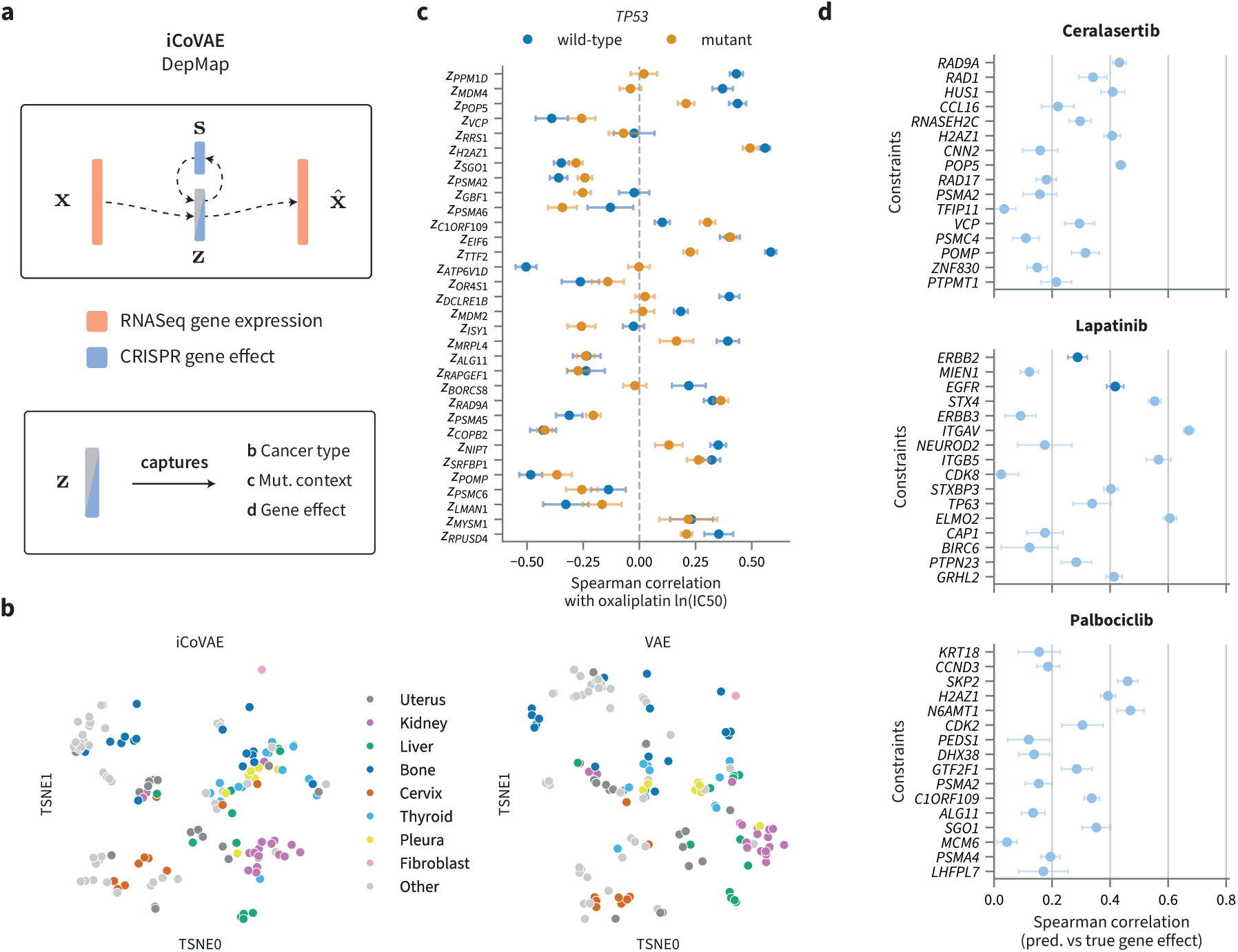
Accuracy of CRISPR gene effect prediction from constrained features and richness of iCoVAE-generated representations. **a** An iCoVAE model is fitted using RNASeq gene expression data from DepMap as an input, and CRISPR gene effect as auxiliary data. The information captured by the generated representations is then studied. **b** t-SNE of iCoVAE-generated representations and VAE-generated representations across a set of cancer types unseen during model training. Points are coloured by cancer type. PNS: peripheral nervous system, UT: urinary tract. **c** Spearman correlation of features in iCoVAE-generated representations with oxaliplatin drug response, for a model constrained by CRISPR gene effect in genes associated with response to oxaliplatin. Correlation shown stratified by *TP53* mutation status. Mean and error bars showing +/- 1 s.d. across 10 random seeds. **d** Performance of CRISPR gene effect prediction from constrained representation features in cancer types unseen during training, using iCoVAE models with representations constrained using CRISPR gene effect in genes associated with response to ceralasertib, lapatinib, and palbociclib, respectively. Points corresponding to the prediction of CRISPR gene effect in drug targets are coloured blue. Mean and error bars showing +/- 1s.d. across 10 random seeds.

To test the ability of constrained dimensions in iCoVAE representations to capture information about the auxiliary data, we looked at how well the constraints themselves (i.e. CRISPR gene effect for the selected genes) are predicted from the constrained dimensions in the representations. We see that iCoVAE is able to effectively predict CRISPR gene effect in samples of cancer types unseen during training, using gene expression data. We draw particular attention to the accuracy of gene effect prediction for the nominal targets of lapatinib (*ERBB2, ρ* = 0.288, *EGFR, ρ* = 0.418 Spearman correlation. Prediction of *ERBB2* gene effect is higher in cross-validation, as very few samples in the test set are dependent on *ERBB2*).

Representations generated by iCoVAE also retain other information from gene expression profiles, which is disentangled from information related to the constraints. This can be seen in Fig. 2b. This information is captured primarily in the unconstrained dimensions of the iCoVAE representation, which operate in a similar way to a standard VAE representation, encoding variation in gene expression in a compact or compressed form.

Next, we studied whether known relationships between CRISPR gene effect and drug response are captured in iCoVAE representations.

Constrained dimensions of iCoVAE representations can encode rich information related to gene effect beyond mutation status, which can be seen by studying the correlation of each representation dimension with response to oxaliplatin, stratified by *TP53* mutation status. In Fig. 2c, we show the correlation between both *z*_*PPM*1*D*_ and *z*_*MDM*4_ (the specific dimensions constrained to capture cell line dependency on these genes) and response to oxaliplatin across cell lines in the unseen test set. It has been observed that lack of p53 function is a key mediator of response to platinum-based therapies such as oxaliplatin [32, 33]. Both of these genes act as inhibitors of p53—an effect which is captured in the dimensions linked to their CRISPR gene effect, as shown by the lack of correlation between these features and response to oxaliplatin in *TP* 53-mutant samples (*z*_*PPM*1*D*_ : *ρ* = 0.020 and *z*_*MDM*4_ : *ρ* = *−*0.040, Spearman correlation) compared to in *TP53* -wild-type samples (*z*_*PPM*1*D*_ : *ρ* = 0.360 and *z*_*MDM*4_ : *ρ* = 0.431). In *TP* 53-mutant samples, neither of these genes can actively inhibit the already inactivated *p*53, and thus the products of these genes do not influence the cancers’ resistance to oxaliplatin. Representation dimensions constrained using CRISPR gene effect in these examples thus demonstrate a capture of information richer than mutation status alone—in this case capturing something closer to *p53* activity rather than solely *TP53* mutation status—and are directly associated with drug response. These relationships are consistent across iterations of the same model, whereas the correlation of unconstrained representation dimensions with drug response varies widely, supporting the notion that PiCo can learn stable features which capture meaningful biological information.

### 2.3 PiCo improves robustness and interpretability of drug response prediction in cancer cell lines

In line with our interest in robustness, we focus on tests of out-of-distribution generalisation, demonstrating generally that the addition of relevant biologically-informed constraints in machine learning models for cancer medicine can improve robustness.

First, we evaluated the performance of PiCo in the challenging setting of performing drug response prediction for cancer cell lines of cancer types which are entirely unseen during model training. For these unseen cancer types, PiCo operates in an “out-of-the-box” fashion, with neither any training data from the unseen cancer type nor any model fine-tuning using this data performed. Gene expression profiles of different cancer types have different underlying distributions [34], influenced by cancer type-specific somatic mutations and copy number aberrations, as well as other alterations affecting gene expression, such as promoter methylation. All of these changes can impact CRISPR gene effect, so we consider this a suitable test of out-of-distribution generalisation.

We used data from the Genomics of Drug Sensitivity in Cancer (GDSC) resource, a pan-cancer pharmacological screen on cancer cell lines [28], for drug response data. Many of the same cell lines are screened in DepMap [27], resulting in the availability of both CRISPR gene effect and drug response data for a total of approximately 500 cell lines (depending on the drug).

For a range of 40 drugs, we trained PiCo models with a heterogeneous training set (spanning multiple cancer types, each with more than 40 samples in DepMap, for further details see Fig. B1). A further 16 cancer types, which were entirely unseen, were reserved and used as a test set for assessing the generalisation of the model. We considered two main baselines for comparison: prediction from VAE-generated representations (similar to iCoVAE-generated representations, but with no use of auxiliary data) and prediction of drug response directly from gene expression using a neural network of a similar complexity to the iCoVAE encoder (referred to as “NN”). For the VAE-based method and PiCo, we considered two different linear drug response prediction models during model selection: linear support vector regression (SVR) and ElasticNet. (PiCo-ElasticNet refers to a PiCo model using ElasticNet as its prediction model, for example. See Section 4 for further details).

Comparing across all drugs and random seeds, PiCo performs significantly better than VAE-based models and NN in terms of RMSE (VAE vs. PiCo *p* = 5.51 *×* 10^*−*12^, NN vs. PiCo *p* = 8.83 *×* 10^*−*34^, Wilcoxon signed-rank test) and better than the VAE-based models in terms of Spearman correlation (VAE vs. PiCo *p* = 4.80 *×* 10^*−*8^, NN vs. PiCo *p* = 0.73, Wilcoxon signed-rank test). For many of the small-molecule inhibitors studied, PiCo provides a substantial improvement in out-of-distribution generalisation (Fig. 3b) over baselines. For the ATR inhibitor ceralasertib, PiCo predicts drug response in cell lines of unseen cancer types with considerably higher accuracy than the best baseline (PiCo-ElasticNet: *ρ* = 0.434, VAE-ElasticNet: *ρ* = 0.307, NN: *ρ* = 0.288, Spearman correlation). This trend was reproduced for the IGF1R inhibitor linsitinib (PiCo-ElasticNet: *ρ* = 0.576, VAE-ElasticNet: *ρ* = 0.515, NN: *ρ* = 0.522), the VEGFR inhibitor axitinib (PiCo-ElasticNet: *ρ* = 0.634, VAE-ElasticNet: *ρ* = 0.572, NN: *ρ* = 0.599), and the dual tyrosine kinase inhibitor lapatinib (PiCo-ElasticNet: *ρ* = 0.224, VAE-SVR: *ρ* = 0.170, NN: *ρ* = 0.179). PiCo can also improve generalisation in the prediction of response to non-targeted drugs, such as the antimetabolite methotrexate (PiCo-ElasticNet: *ρ* = 0.394, VAE-ElasticNet: *ρ* = 0.345, NN: *ρ* = 0.345) and for platinum-based chemotherapies such as oxaliplatin (PiCo-ElasticNet: *ρ* = 0.535, VAE-ElasticNet: *ρ* = 0.492, NN: *ρ* = 0.544).

**Fig. 3.**
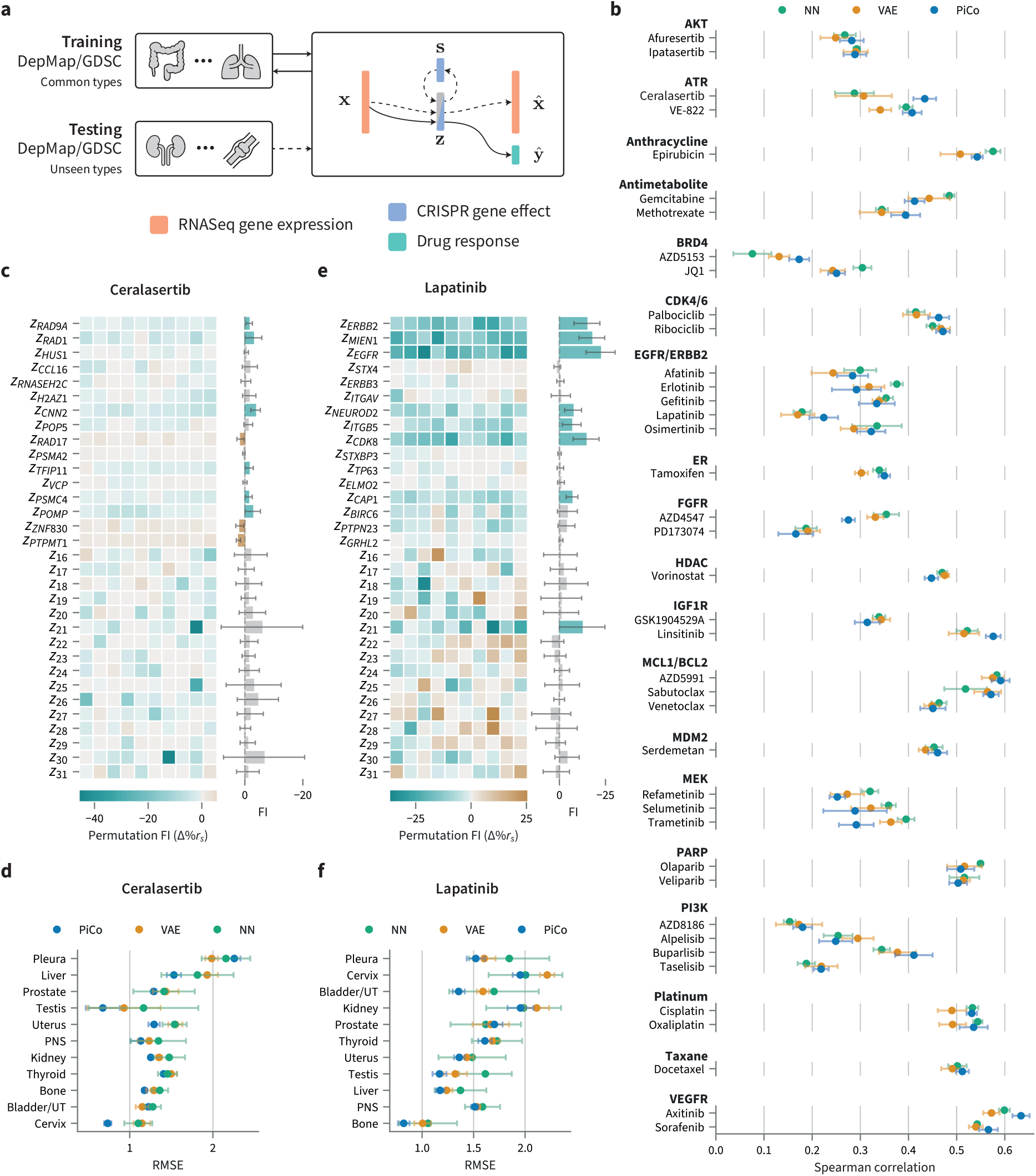
Robustness and feature importance in PiCo. **a** To assess model generalisation to unseen cancer types, feature extractors and prediction models were trained on cell lines of common types, with samples of less common types reserved for testing in zero-shot drug response prediction. **b** Prediction performance in unseen cancer types, aggregated across all types, for a range of drugs. Drugs are grouped by nominal target, or targeted pathway. Mean Spearman correlation and +/- 1 s.d. across 10 random seeds. **c** (left) Permutation feature importance (FI) for the model predicting ceralasertib response in test samples for each feature in PiCo-ElasticNet model, for 10 random seeds. (right) Marginal plot of FI across 10 random seeds. Error bars show +/- 1 s.d. Bars for features where perturbation consistently decreases performance are coloured blue. Bars are coloured grey if the mean FI +/- 1 s.d. includes zero FI. **d** Prediction performance in unseen cancer types, separated by cancer type. Spearman correlation. **e** and **f** as in **c** and **d**, for lapatinib.

Across the small molecule inhibitors for which PiCo provides a benefit in generalisation for drug response prediction, PiCo prediction models inform their predictions of drug response using representation dimensions constrained using CRISPR gene effects of the given drugs’ nominal targets. Fig. 3c and Fig. 3d demonstrate this effect using permutation-based feature importances from PiCo drug response prediction models (i.e. the linear model trained to predict drug response from generated representations. See Section 4 for details). For lapatinib, the importance of the representation dimensions constrained using CRISPR gene effect of its nominal targets *EGFR* and *ERBB2* are high, and this feature importance is stable across multiple random seeds (*EGFR*: *FI* = *−*22.1%, *ERBB2* : *FI* = *−*14.7% reduction in Spearman correlation after random feature permutation) (Fig. 3e). For unconstrained representation dimensions, there is a large variation in feature importance, since these dimensions do not necessarily capture the same information in each instance of the same model, unlike those which are constrained. For some targeted therapies, dimensions constrained to capture CRISPR gene effect of genes encoding proteins which interact with the drug’s nominal target are also key: in prediction of response to ceralasertib, an ATR inhibitor, consistently important dimensions include features constrained by *RAD1* and *RAD9A* CRISPR gene effect (*RAD1* : *FI* = *−*3.01%, *RAD9A*: *FI* = *−*1.51%) (Fig. 3c). *RAD9A, RAD1* and *HUS1* form a complex which is required to trigger ATR-mediated Chk1 activation for DNA damage repair [35].

For oxaliplatin, a platinum-based chemotherapy, we see high importance assigned to dimensions constrained using *MDM4* and *TTF2* CRISPR gene effect (*FI* = *−*8.02%, *FI* = *−*5.55%, respectively, Fig. B2. Although the products of these genes are not directly targeted by oxaliplatin, there are plausible biological explanations for why a cancer cell line’s dependence on the function of each of these genes may be indicative of its response to the drug. Inactivation of p53 is a common factor in resistance to oxaliplatin [33], and wild-type p53 is often inactivated through high expression of *MDM4* [32]. A dependency on *MDM4* suggests a reliance on its function as an inhibitor of p53 to induce proliferation. For a second commonly used platinum-based chemotherapy, cisplatin, we see high importance assigned to features constrained using *RAD9A* and *TXNRD1* CRISPR gene effect (*FI* = *−*2.43%, *FI* = *−*7.05%, respectively, Fig. B2). Both of these genes are known to be involved in response to cisplatin—*RAD9A* through its role in DNA damage repair [36] and *TXNRD1* through its role in response to oxidative stress [37]. For the taxane, docetaxel, the dimension constrained using *MCL1* CRISPR gene effect is highly influential in its predictions on unseen cancer types (*FI* = *−*5.48%, Fig. B2). *MCL1* is known to regulate response to taxanes [38].

The performance gains that PiCo offers vary across the set of unseen cancer types and broadly reflects the relevance of drugs and their targets to given cancer types. For example, we see a large benefit provided by PiCo for prediction in unseen cell lines derived from cancers of the uterus and cervix in prediction of response to ceralasertib (Uterus: PiCo-ElasticNet: RMSE = 1.287, *ρ* = 0.282, VAE-ElasticNet: RMSE = 1.494, *ρ* = *−*0.143, NN: RMSE = 1.540, *ρ* = *−*0.167, cervix: PiCo-ElasticNet: RMSE = 0.735, *ρ* = 0.668, VAE-ElasticNet: RMSE = 1.043, *ρ* = 0.190, NN:RMSE = 1.103, *ρ* = 0.253, Fig. 3)—inhibiton of ATR is particularly promising in gynaecological cancers due to the prevalence of homologous recombination deficiency (HRD) in these cancers which leads to a reliance on ATR for DNA damage repair (DDR). This reliance on ATR is particularly strong in cervical cancers, likely due to the link between human papilloma virus (HPV)—the primary cause of cervical cancers—and DDR pathways. Similar patterns demonstrating the specificity of PiCo performance gains to relevant cancer types are seen for other drugs, both targeted and non-targeted (Section A.2).

The general performance trend in the test set is reproduced in cross-validation (i.e. using data from all but the 16 least represented cancer types in DepMap), with PiCo performing significantly better than the VAE-based model in terms of RMSE (VAE vs. PiCo *p* = 6.64 *×* 10^*−*11^, Wilcoxon signed-rank test) and Spearman correlation (VAE vs. PiCo Wilcoxon signed-rank test *p* = 1.74 *×* 10^*−*6^). Extending feature importance analysis into results from the cross-validation process further demonstrates the ability of PiCo to base predictions on biologically meaningful features. For the previous examples, ceralasertib and lapatinib, features important for prediction in the held-out cancer types are similarly important in cross-validation (Fig. B3). This suggests that these features capture information which is predictive of drug response independently of cancer type.

Of further interest are the drugs for which PiCo performs *worse* than baselines on the held-out cancer types. The most extreme example is for the MEK inhibitor trametinib (PiCo-SVR: *ρ* = 0.292, VAE-ElasticNet: *ρ* = 0.363, NN: *ρ* = 0.395, Fig. 3). When instead considering performance in the cross-validation procedure, PiCo performs substantially better than the same baselines (PiCo-SVR: *ρ* = 0.730, VAE-ElasticNet: *ρ* = 0.686, Fig. B3). By comparing feature importances in the held-out cancer types and the cancer types involved in the cross-validation process, we can explain why this difference in performance occurs. Features encoding *BRAF, CTNNB1, ZEB2* and *RPS2* CRISPR gene effect are highly important in the validation sets, but reliance on these features negatively impacts performance in the held-out set. This suggests that the association between these features and trametinib response is context-specific, or that these dependencies are difficult to predict in the new cell lines. Some of these features are dependencies that are much more common in certain cancer types, such as *BRAF* and *ZEB2* in melanoma.

### 2.4 PiCo for clinical data

Next, we sought to investigate whether the proposed models could also be useful for clinical prediction. We applied PiCo in three clinical breast cancer cohorts, TransNEO [29], SCAN-B [30], and The Cancer Genome Atlas (TCGA) [31], first fitting iCoVAE using only cell line data, then generating representations of unseen patient gene expression data, which are used in PiCo.

#### 2.4.1 iCoVAE features capture pathway interactions and are associated with outcomes in patients while preserving subtype-specific features

For each clinical dataset, we began by fitting an iCoVAE model using cell line data with constraints related to four drugs, which comprise the common breast cancer chemotherapy treatment regimen, FEC-T (5-fluorouracil, epirubicin, cyclophosphamide, and a taxane, either paclitaxel or docetaxel)(See Methods for details). This treatment regime is used in the TransNEO cohort [29]. Patients with ERBB2 amplified breast cancer in the TransNEO cohort are also treated with trastuzumab, an anti-HER2 antibody. Since no such screen using trastuzumab exists in cancer cell lines, we add constraints related to gene effect in genes known to influence anti-HER2 therapy sensitivity: *ERBB2, ERBB3, PIK3CA* and *EGFR*. Representations of patient gene expression data from the TransNEO cohort and its external validation set ARTemis+PBCP were generated, as were representations of patient gene expression in the SCAN-B and TCGA cohorts, with TCGA used as an external validation set for a model fitted using SCAN-B data. All representations were generated using the pretrained cell line model.

In Fig. 4c and Fig. 5c, we see that representations of patient gene expression data generated by iCoVAE preserve differences across breast cancer molecular subtypes in patients, despite being fitted on a dataset of pan-cancer cell lines where less than 5% are derived from breast cancers. In both datasets, samples clearly group by molecular subtype, with a clear separation of triple-negative breast cancers (TNBC).

**Fig. 4.**
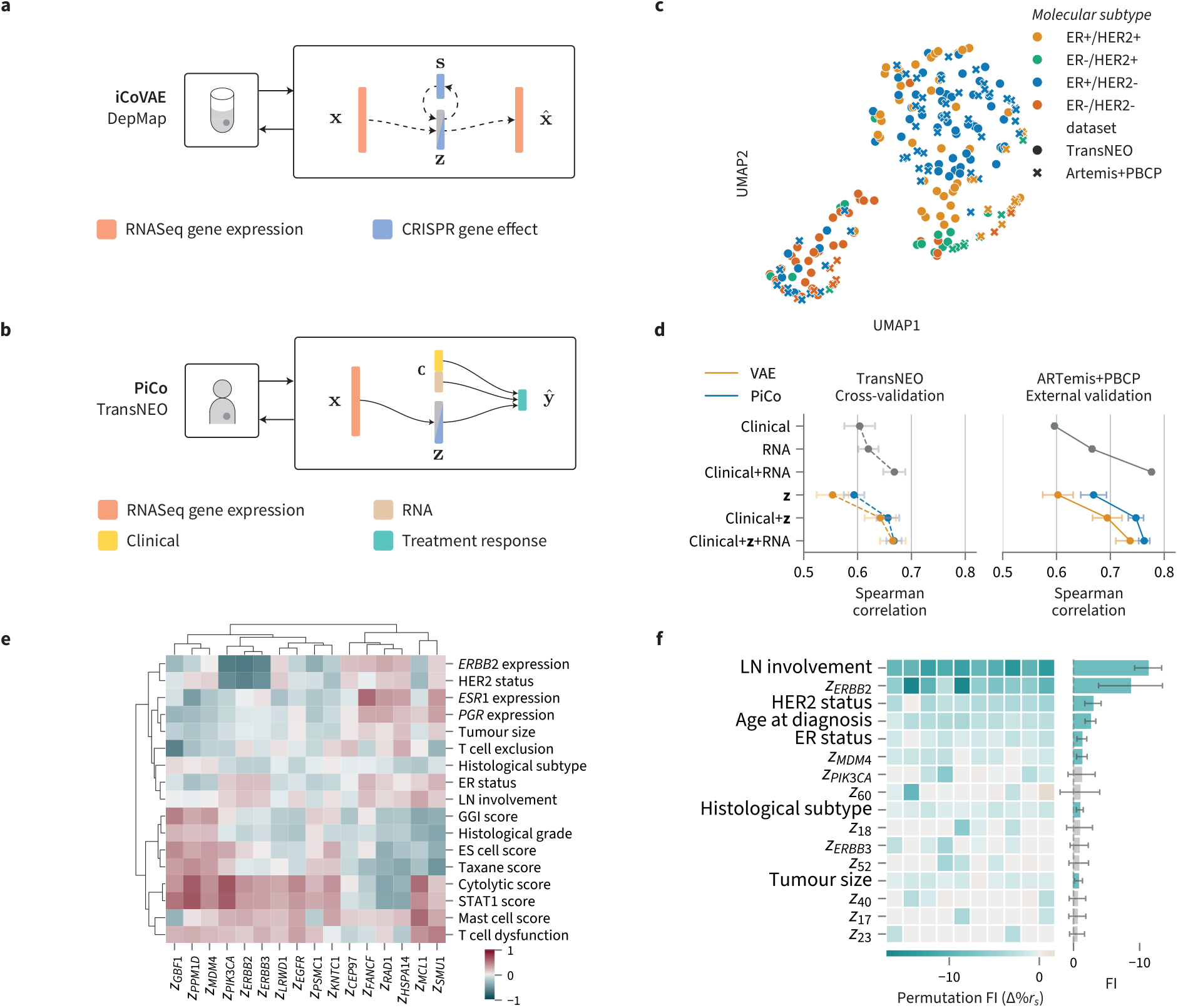
PiCo for predicting neoadjuvant treatment response in breast cancer patients from the TransNEO cohort. **a** An iCoVAE model is fitted to capture information about CRISPR gene effect in cell lines from the DepMap dataset, using gene expression data as an input. **b** The encoder from the fitted iCoVAE is used to generate representations for patient gene expression data. These representations are then combined with clinical features and hand-engineered features based on gene expression to predict treatment response. **c** UMAP of patients in training and external validation cohorts based on representations generated by iCoVAE. **d** Prediction of residual cancer burden (RCB) score in TransNEO, in cross- validation and the ARTemis+PBCP external validation set. 5-fold cross-validation in 10 random seeds. Error bars display +/- 1 s.d. around the mean across 10 seeds. Models are named according to the inputs to regression models. **e** Pearson correlation between iCoVAE features and other clinical and hand-engineered features available in TransNEO. **f** Permutation feature importance for regression models predicting RCB score. Δ%*ρ*_*s*_ is the percentage change in Spearman correlation after random shuffling of the given feature. Features with the 15 largest negative Δ%*ρ*_*s*_ values are shown. Negative values indicate Spearman correlation decreases upon perturbing the feature. Δ%*ρ*_*s*_ shown separately across 10 random seeds and aggregated, with error bars showing +/- 1 s.d. around the mean. Clinical+**z** models shown.

**Fig. 5.**
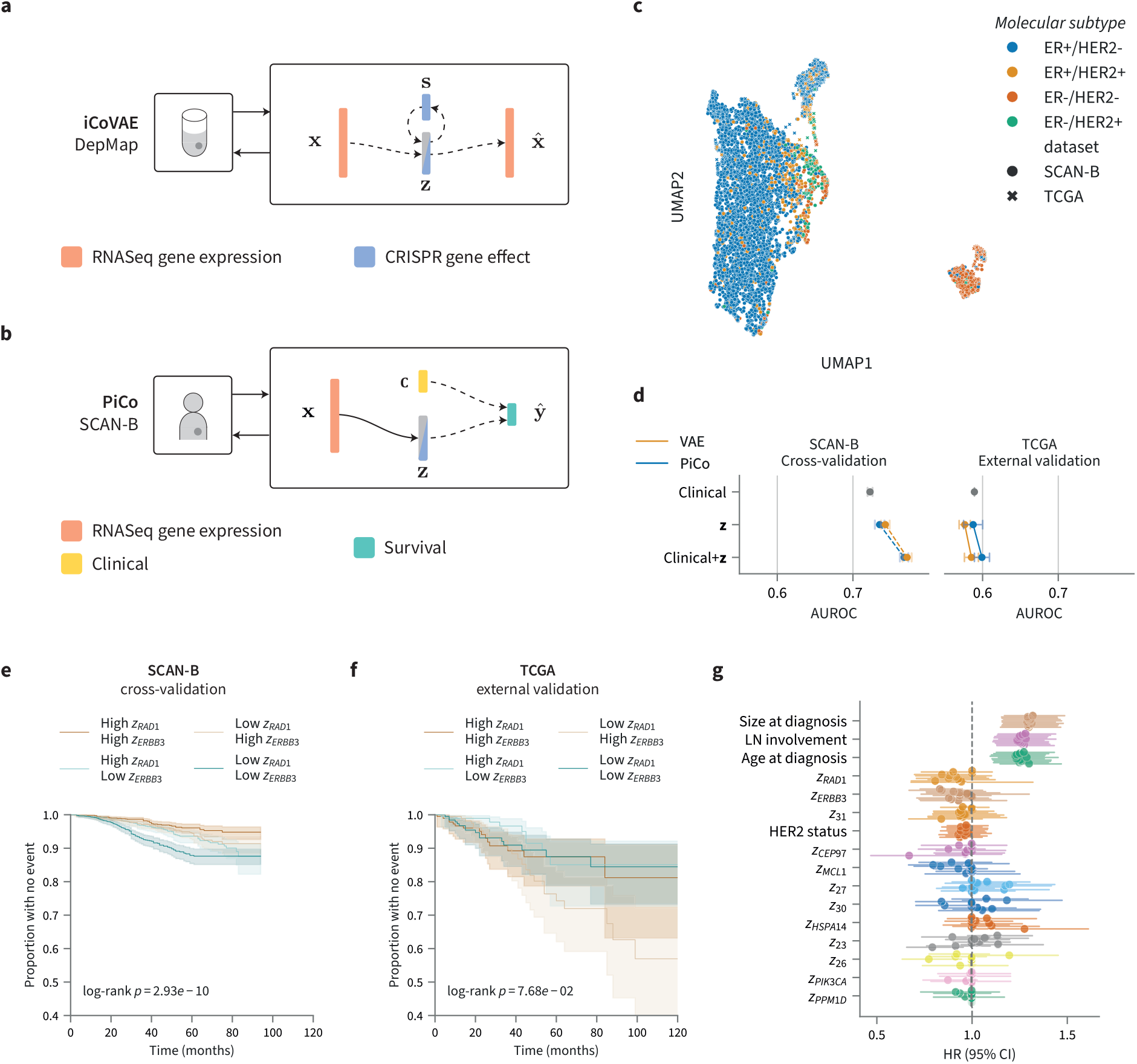
PiCo for breast cancer free survival (BCFS) in breast cancer patients from the SCAN-B cohort. **a** An iCoVAE model is fitted to capture information about CRISPR gene effect in cell lines from the DepMap dataset, using gene expression data as an input. **b** The encoder from the fitted iCoVAE is used to generate representations for patient gene expression data. These representations are then combined with clinical features to predict BCFS. **c** UMAP of patients in training and external validation cohorts based on representations generated by iCoVAE. **d** Prediction by logistic regression of 5-year breast cancer free survival (BCFS), using AUROC as a metric. 5-fold cross-validation in 10 random seeds. Error bars display +/- 1 s.d. around the mean across 10 seeds. Models are named according to the inputs to the models. **e** Kaplan- Meier plot showing difference in survival for SCAN-B patients with high and low *z*_*RAD*1_ and *z*_*ERBB*3_. Patients split above and below the median for each feature, random seed 30. Bands show 95% confidence interval. **f** As in **e** for TCGA external validation. **g** Hazard ratios for PiCo Clinical+**z** Cox model (stratified by ER status). Hazard ratios and 95% CI shown for 10 random seeds. Sorted by median hazard ratio difference from 1.

Next, we sought to understand whether features from representations of patient gene expression data generated by iCoVAE capture known relationships between clinically relevant features. Fig. 4e displays the association of features in representations generated by iCoVAE for patient gene expression data with other clinically-relevant features available for TransNEO (an equivalent plot for SCAN-B is given in Fig. B6). *z*_*PPM*1*D*_ is strongly correlated with *ESR1* expression in TransNEO (*ρ* = *−*0.483, Fig. 4e). The link between Wip1, the protein encoded by *PPM* 1*D*, and ER*α* has been studied extensively, evidencing a positive feedback loop between Wip1 and ER*α* which increases proliferative signalling in cancers with functional p53 [39].

Finally, we studied the association of individual features extracted by these models with outcomes in each of these datasets: in TransNEO, we considered association with residual cancer burden (RCB) score, a score which considers primary tumour size and lymph node positivity after neoadjuvant therapy. In SCAN-B, we considered association with 5-year breast cancer-free survival (BCFS). *z*_*PPM*1*D*_ exhibits the strongest association with RCB score of the features generated by iCoVAE (*ρ* = *−*0.438)—an association which is reflected in the external validation dataset ARTemis+PBCP (*ρ* = *−*0.339). In SCAN-B, lymph node status is most predictive of 5-year breast cancer free survival (BCFS) (AUROC = 0.563), followed closely by *z*_*MCL*1_, HER2 status, and tumour size at diagnosis (AUROC = 0.558, AUROC = 0.550 and AUROC = 0.549, respectively).

To identify features which are predictive of outcome in specific cancer subtypes, we also considered associations of features extracted by iCoVAE models with outcome for each breast cancer molecular subtype separately in each cohort (Fig. B5 and Fig. B6). As expected, in the TransNEO cohort, there is a strong association between *z*_*PIK*3*CA*_ and *z*_*ERBB*2_ and treatment response in patients with ER+/HER2+ breast cancers, and an association between *z*_*PPM*1*D*_ and treatment response in ER+/HER2-cancers. In the latter case, this association is even stronger than the association between *ESR*1 expression and treatment response.

#### 2.4.2 Features linked to gene dependency are robust predictors of treatment response in patients

For the TransNEO cohort, we trained models to predict residual cancer burden (RCB) score using an ElasticNet-regularised regression in the PiCo framework, using 10 separate 5-fold cross-validations to optimise hyperparameters (see Methods). The hyperparameters producing the best performance across cross-validations for each of the 10 random seeds were then used to refit the models on the entire TransNEO dataset. Model performance across these 10 models was then evaluated on ARTemis+PBCP, an external validation set. We included models using various combinations of modalities as inputs: Clinical features, human-engineered features derived from RNASeq gene expression data, including features related to the tumour immune microenvironment, as used in [29] (referred to as RNA), and representations generated by iCoVAE and VAE models (referred to as **z**). We then further consider using predictions of RCB score as predictions of pathological complete response (pCR), defined as an RCB score of zero.

In TransNEO, the performance of PiCo and the VAE-based model in their prediction of RCB score in both TransNEO and ARTemis+PBCP is strong (Fig. 5d). The addition of constrained features to representations in the PiCo Clinical+**z** model provides a performance benefit on the external validation set over the model using VAE-generated representations (VAE Clinical+**z**: *ρ* = 0.694, PiCo Clinical+**z**: *ρ* = 0.747, Clinical+RNA: *ρ* = 0.776, Spearman correlation, Fig. 5d) and in cross-validation (VAE Clinical+**z**: *ρ* = 0.642, PiCo Clinical+**z**: *ρ* = 0.656, Clinical+RNA: *ρ* = 0.668, Spearman correlation, Fig. 5d). The PiCo Clinical+**z** model is also a clear improvement over the Clinical model (Clinical: *ρ* = 0.596). Models incorporating representations learned by iCoVAE all exhibit improved performance in prediction of pathological complete response (pCR) beyond all other models in the external validation set (PiCo Clinical+**z**+RNA: AUROC= 0.901, VAE Clinical+**z**+RNA: AUROC= 0.878, Clinical+RNA: AUROC= 0.879, Fig. B5). These results demonstrate the general utility of low-dimensional representations of gene expression data in clinical prediction alongside human-engineered features such as genomic grade index (GGI) and features based on single-sample gene set enrichment analysis (ssGSEA).

In addition to producing more robust predictions than other models, PiCo provides interpretability and the potential for identification of mechanisms of response to therapy, whereas models based on VAE representations are more difficult to interpret. Fig. 5f shows feature importances for PiCo models predicting RCB score, using clinical and representation features (Clinical+**z**). Multiple constrained features in PiCo are consistently beneficial to model performance, which is in agreement with their previously discussed association with RCB score. For the Clinical+**z** model, features linked to *ERBB*2 and *MDM* 4 gene effect have the highest feature importance in the external validation set, with a feature linked to *PIK3CA* gene effect close behind (FI= *−*8.726% and FI= *−*1.276%, FI= *−*1.143% respectively, Fig. 4f).

#### 2.4.3 Features linked to gene dependency are useful prognosticators in patients

In SCAN-B, we used a penalised logistic regression as a prediction head within the PiCo framework to predict 5-year breast cancer-free survival (BCFS) in patients. Although SCAN-B covers a range of patients receiving both neoadjuvant and adjuvant therapies, endocrine therapy and immunotherapy, we generated representations from the same iCoVAE models as for TransNEO to study transferability across clinical applications. We similarly carried out 10 separate 5-fold cross-validations to optimise hyperparameters for the Cox proportional hazards regression, splitting the SCAN-B dataset into training and validation datasets, and then reporting the average performance in cross-validation and in the external validation dataset generated from TCGA. This external validation set represents an OOD generalisation test since the TCGA cohort generally consists of younger patients with more advanced tumours, and as such, the rate of events is higher. The age of patients in SCAN-B is generally higher, and these patients tend to have earlier-stage breast cancers (Fig. B7). These differences greatly influence treatment decisions and surgical intervention, as well as outcomes.

Using PiCo, the prediction of 5-year BCFS is improved by utilising representations generated by iCoVAE and VAE models, with synergy between clinical features and representations observed. PiCo performance is similar to using VAE representations in cross-validation (VAE: **z** AUROC = 0.743, Clinical+**z** AUROC = 0.772, PiCo: **z** AUROC = 0.735, Clinical+**z** AUROC = 0.768, Fig. 5c), but improved in external validation (VAE: **z** AUROC = 0.576, Clinical+**z** AUROC = 0.585, PiCo: **z** AUROC = 0.588, Clinical+**z** AUROC = 0.599).

When interrogating feature importance for the PiCo Clinical+**z** model in predictions on the test set (Fig. 5e), we saw high importance assigned to tumour size and lymph node status (FI = *−*0.013, FI = *−*0.008, ΔAUROC), as well as features from the representation such as features linked to dependency on *ERBB2, CEP97* and *MDM4* (FI = *−*0.014, FI = *−*0.008, FI = *−*0.007, Fig. B6).

We finally used features from iCoVAE representations as inputs to penalised Cox regression models (stratified by ER status), to predict the risk of event in breast cancer-free survival. The PiCo Clinical+**z** model provides a better prediction of risk than the Clinical model in a 5-fold cross-validation (c-index = 0.756 and c-index = 0.677, respectively, mean across 10 random seeds, Fig. B6). For *z*_*RAD*1_ in the Clinical+**z** model, the mean hazard ratio HR = 0.910 across 10 seeds. This feature is significant for a single seed, and borderline significant otherwise. *z*_*ERBB*3_ in the Clinical+**z** model, mean HR = 0.912, is also significant in a single seed. To further investigate the effect of these features on BCFS, we separated patients in SCAN-B into groups of high and low *z*_*RAD*1_ and *z*_*ERBB*3_ (above and below the median) and compared their survival curves. Patients stratified into these groups exhibited consistently significantly different risk of event in SCAN-B (min. *p* = 3.39 *×* 10^*−*15^, max. *p* = 6.38 *×* 10^*−*4^, log-rank test, across 10 random seeds, seed 30 shown in Fig. 5), but not always in TCGA (min. *p* = 1.49 *×* 10^*−*2^, max. *p* = 0.81 across 10 random seeds).

## 3 Discussion

In this work, we presented an approach to deep representation learning for omics data that leverages auxiliary data from genetic loss-of-function screens to improve robustness and interpretability. We showed that our model, PiCo, is capable of broad generalisation, including to entirely unseen cancer types and from cell lines to clinical data, whilst retaining a high degree of interpretability.

The idea of placing constraints on the latent space in variational autoencoder-based models has been approached in some recent works by making modifications to the prior over the latent variables. Constraining latent features to capture the mean gene expression of a gene set in cancer, and separating VAE latent spaces into subspaces which capture technical and biological variation separately in single-cell atlases, have been considered [40, 41]. The former utilises prior knowledge from curated pathways, similarly to previously mentioned works, while the latter uses covariates as input data, requiring biological and technical covariates at test time when generating representations for new samples (i.e. fully supervised). Our model differs from these in that it learns from auxiliary data to constrain latent spaces in a semi-supervised way, extracting improved representations of gene expression data without requiring any auxiliary data at test time. A similar idea has recently been proposed in a semi-supervised factor analysis framework, but with the different intention of improving new factor discovery by removing known sources of variation [42]. Although we focused on the use of RNASeq gene expression data, it would also be possible to fit iCoVAE using gene expression data generated from microarrays, or using other omics data types such as copy number, methylation or mutation data, or to extend the model to a multi-omic setting. To further pretrain the iCoVAE, it would also be possible to add further RNASeq data to the unsupervised set of samples used in iCoVAE pretraining from heterogeneous sources such as clinical samples from PCAWG [43]—our models could be used as a starting point for larger models for pre-training on very large gene expression datasets with an eye towards taskand possibly disease-agnostic modelling.

By using a pretrained encoder to generate embeddings and simple linear models to predict targets, we demonstrate the lightweight out-of-the-box utility of the PiCo framework—representations can be generated for any gene expression profile at a very low computational cost, and any model with any level of complexity can be fitted using these representations as inputs. Our results on the use of iCoVAE across different settings support the notion that these constrained models are transportable across cancer types in cell lines, and between cell lines and patient tumours. This is possible despite the fact that there are large differences in gene expression profiles between these settings.

In general, the discovery and validation of robust signatures for prognosis and prediction of treatment response based on transcriptomic data is a difficult task due to experimental variability and complex correlation patterns across the transcriptome. Many simple and thus interpretable (e.g. mean expression in a handful of genes) gene expression-based predictors of treatment response or prognosis have been developed [44–47] and some have been used in models for prediction of clinical outcomes [29]. Some of these signatures, such as the paclitaxel response metagene [47] were developed with the aid of loss-of-function screening in cancer cell lines. We have shown that more complex but similarly interpretable features extracted by an entirely data-driven model fitted on cell lines can provide a similar level of indication of patient response to treatment as manually-engineered scores developed with high levels of domain expertise. We therefore believe that the further development and validation of these interpretable but complex, data-driven indicators of therapy response presents a potential for clinical impact in cancer medicine.

While PiCo can provide a benefit over compared baselines in the settings studied, we observed varying levels of improvement across cancer types and drug targets. In cases where PiCo does not provide benefit, this is mostly due to the lack of generalisation of a feature linked to gene effect to the target setting, either because of a change in the relationship between gene effect and drug response across settings, or because of a failure in the prediction of gene effect in a new setting. This demonstrates that the constraint selection step of PiCo is extremely important for influencing its out-of-distribution generalisation. This step is performed separately to the drug response prediction stage of model training, and therefore, it is not possible to know the optimal selection of gene effects to which dimensions of iCoVAE should be linked for a given task. This selection is also required to be different for each drug considered in the current formulation, meaning iCoVAE representations do not necessarily generalise across drugs, unless selected constraints are similar. Although this can be the case for small-molecule inhibitors with the same nominal target, no general utility iCoVAE representation is currently generated. The use of a limited number of linked dimensions, constrained by the size of the latent space of the iCoVAE, also presents another limitation. A direction for the future development of the PiCo framework for drug response prediction could include the development of a general utility iCoVAE integrating many gene effects, and a PiCo model which extends to multi-drug prediction by incorporating drug structural information as an input.

While we focused on the zero-shot prediction setting for new cancer types, few-shot learning has also shown promise on similar tasks [48]. The PiCo framework can be naturally extended to this setting, allowing fine-tuning of drug response prediction heads and the iCoVAE encoder itself, although care would need to be taken to preserve the meaning of constrained dimensions and therefore their interpretation [49].

The PiCo framework could also be extended to other settings outside of single drug response prediction, such as the prediction of cell line response to combinatorial CRISPR knockouts or combination drug treatments, or to applications outside of cancer medicine entirely. Our method is generally applicable to any two related datasets owing to the two-stage training process. In any setting where transfer learning could be reasonably applied (i.e. there is a meaningful link between the two datasets) PiCo can be used to do this in an interpretable way.

## 4 Methods

### 4.1 Model design

We aim to use information from DepMap, a large-scale genetic loss-of-function screen on cancer cell lines, to improve downstream models in a variety of scenarios, inferring links between dependency of a cancer on gene function and outcomes such as drug response in cell lines, as well as outcomes in patients such as treatment response and disease-free survival. In the DepMap study, a CRISPR-Cas9 system is used to individually knock out around 17,000 genes across around 1000 cell lines. Many cell lines have also undergone extensive molecular characterisation through the Cancer Cell Line Encyclopedia (CCLE) project, including RNASeq gene expression.

#### 4.1.1 VAE

In a vanilla VAE, low-dimensional representations of observations **x** ∈ ℝ ^*d*^ are learned through variational inference, linking observations to a latent variable **z** ∈ ℝ ^*L*^. The likelihood of an observation **x** is factorised as:

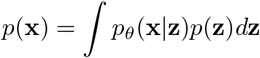

The aim is to learn the parameters of the distribution *p*_*θ*_(**x**|**z**), which is approximated by a neural network with parameters *θ*. This distribution is typically referred to as a decoder. An amortized variational approximation to the intractable posterior *p*(**z**|**x**) is also learned, which is referred to as *q*_*ϕ*_(**z**|**x**) and is also approximated by a neural network with parameters *ϕ*. This distribution is typically referred to as an encoder.

We then define a lower bound on the log-likelihood *p*(**x**) to be used as a model objective:

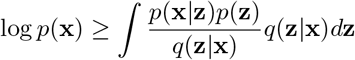

This is then optimised as:

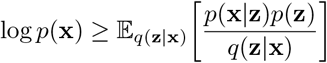

In our particular implementation, we use a calibrated decoder as in our proposed model to ensure a fair comparison.

#### 4.1.2 iCoVAE and PiCo

For representation learning of gene expression data, we developed a semi-supervised variational autoencoder-based [13] model suitable for utilising CRISPR gene effect data as auxiliary information. We refer to this model as interpretable constrained VAE (iCoVAE).

Supervision in the iCoVAE comes in the form of a conditional prior on a subspace of the latent space, as well as a regression model in which multiple univariable linear models predict gene effect for a selected set of genes from individual features in the latent space. Thus, the encoder is trained to encode features in latent representations that are linear predictors of gene effect. This linear dependence allows for interpretability when representations are used in downstream models. In addition to this, the model preserves additional information encoded in gene expression profiles through the unconstrained features in the representation, which are not linked to any gene effect.

We assume the availability of a dataset 𝒟 = 𝒟_*x*_ ∪ 𝒟_*x,s*_ ∪ 𝒟_*x,s,y*_ ∪ 𝒟_*x,y*_ where 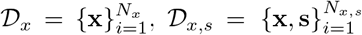 and so on. *X*^(*s*)^ as in Fig. 1 then refers to the matrix formed by stacking data points from 𝒟_*x,s*_ and 𝒟_*x,s,y*_ (for details on data splitting see Section 4.7). The iCoVAE builds both upon the semi-supervised variational autoencoder framework [13] and the Characteristic Capturing VAE (CCVAE) [10], 2 with likelihood of some observed data, drawn from 𝒟, consisting of **x** ∈ ℝ^*d*^ and/or added auxiliary information for supervision 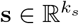 taking the form:

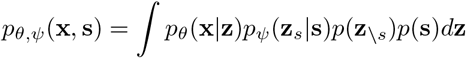

where in this particular application, **x** is a random vector representing gene expression, **s** is a random vector capturing the gene effects of the chosen genes for the model, **z** ∈ R^*L*^ is the latent vector, which is decomposed into *k*_*s*_ dimensions linked to gene effect in **z**_*s*_ and *k*_*\s*_ unlinked dimensions in **z**_*\s*_. The output distribution *p*_*θ*_(**x**|**z**) is parameterised by a neural network with parameters *θ* as 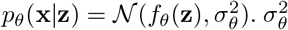 is a shared parameter which improves decoder calibration when dimensions in **x** have differing variances and ease of reconstruction [50]. The conditional prior *p* (**z** |**s**) factorises as 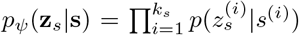 for the linked features of the latent space and takes the form *p*(**z**_*\s*_) = *𝒩* (**0, I**) for the unconstrained features. The distribution for each individual latent dimension in **z**_*s*_ is further defined as 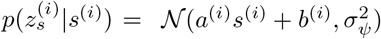 where the parameters *a*^(*i*)^, *b*^(*i*)^, and 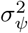 are learned.

To learn the parameters of these distributions, we utilise variational inference, modelling an amortised variational approximation to the true, intractable posterior *p*(**z**|**x, s**), which we refer to as *q*_*φ,ϕ*_(**z**|**x, s**). Rather than modelling this distribution directly, it is factorised as

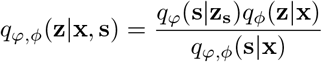

where *q*_*φ,ϕ*_(**s**|**x**) = ∫*q*_*φ*_(**s**|**z**_**s**_)*q*_*ϕ*_(**z**|**x**)*d***z**.

The distribution of the regression model *q*_*φ*_(**s**|**z**_**s**_) is defined similarly to the conditional prior, such that 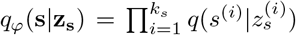. The predictive distribution for each gene effect is defined as 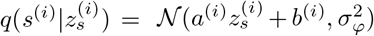 where the parameters *a* ^*(i)*^, *b* ^*(i)*^, and 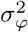 are learned. 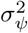 and 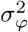 are learned as shared variances for the regressor and conditional prior respectively, in a similar fashion to the shared variance 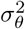 which is learned for the decoder (more detail on this choice is available in Supplementary Information).

To define a model objective, we define a lower bound on the log-likelihood

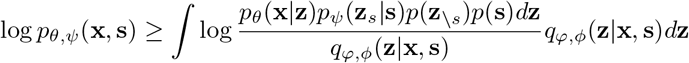

With the term *q*_*φ,ϕ*_(**z**|**x, s**) defined as above.

The lower bound on the log-likelihood log *p*_*θ,ψ*_(**x, s**) can be written and optimised as

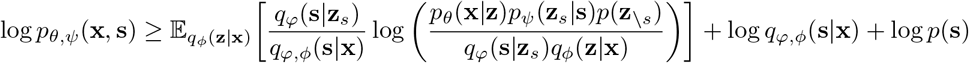

For cell lines with gene expression data but no CRISPR screening data, we can perform variational inference in a similar sense, treating **s** as a latent variable which can be marginalised over, obtaining

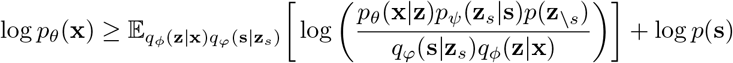

It is worth noting that **s** can be any data type, given an appropriate distribution *q*_*φ*_(**s**|**z**_*s*_) is used. Fitting the iCoVAE defines the first stage of the PiCo framework. In the second stage, PiCo employs a *linear probing* method to predict a target from representations generated by a frozen, pretrained iCoVAE encoder (Fig. 1). The iCoVAE encoder is used to generate representations using fixed parameters learned during the previous stage of modelling, and a linear prediction head is fitted to map from the representation to the target. For a given new data point **x**^∗^, the maximum a posteriori (MAP) representation, taking **z**^∗^ = arg max *q*_*ϕ*_(**z**|**x**^∗^), is used, which provides a balance between performance in downstream tasks and computational efficiency [51].

The choice of linear probing is motivated by the desire for performance in an out-of-distribution setting and for preserving model interpretability by avoiding fine-tuning [49]. In this particular implementation, the prediction head is either a support vector regression with a linear kernel (SVR) or an ElasticNet regularised linear regression.

This method for prediction, which combines an iCoVAE encoder with a linear prediction head, forms the proposed PiCo model.

### 4.2 Constraint selection

The relationship between drug responses and the gene effect of their nominal targets and neighbours in gene functional association networks has been studied in the context of mechanism-of-action discovery [52]. For our application, we are interested in predicting gene effects, which are in turn most predictive of drug response. Since we aim to predict drug response from representations with a linear model, we aim to select the most robust *linear* predictors of drug response from the set of gene effects studied in DepMap. Previous studies of the relationship between drug response and gene effect datasets have removed principal components from the drug response dataset which are correlated with cell line growth rate, such that these do not interfere with the discovery of mechanism of action. Since we are concerned with predicting drug response, inclusive of the effects of cell line growth rate on response, we choose to preserve these components.

For each candidate constraint, we compared two linear models, similarly to [52]. The first model, representing the null hypothesis, predicts drug response using only CRISPR screen experimental confounders as covariates: (i) growth pattern (suspension, adherent, or semi-adherent), (ii) median depletion caused by knockout of essential genes, (iii) screen ROCAUC (defined as the area under the receiver operating characteristic curve for essential versus non-essential control genes), (iv) mean absolute deviation of depletion for essential gene knockouts, and (v) tissue of origin of the cell line. These confounders are chosen due to their association with gene effect across many knockouts, particularly pan-essential genes.

The alternative hypothesis includes the covariates of the null model plus the gene effect of a single gene. The models are defined as follows:

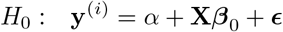

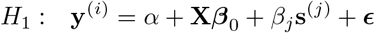

where **y**^(*i*)^ ∈ ℝ ^*n*^ is the vector of drug response values (log-transformed IC50) across *n* cell lines for drug *i*, **X** ∈ ℝ ^*n×p*^ is the matrix of confounders, ***β***_0_ ∈ ℝ ^*p*^ is the vector of regression coefficients for the confounders, **s**^(*j*)^ ∈ ℝ^*n*^ is the vector of gene effects for gene *j*, and *β*_*j*_ ∈ R is its associated coefficient. The scalar *α* is an intercept term, and ***ϵ*** is an error term.

The significance of each gene effect in predicting response to a drug is assessed using a likelihood ratio 3 test comparing the alternative model *M*_1_ to the null model *M*_0_. The test statistic is:

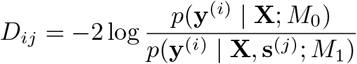

Under the null hypothesis, *D*_*ij*_ ∼ *χ*^2^(1), with 1 degree of freedom.

We repeat this procedure for each of the 17,931 genes screened in DepMap, for each drug. Benjamini-Hochberg false-discovery rate (FDR) correction is performed per drug, rather than across all drugs. This choice is made to account for differences in strengths of associations between drugs—for targeted drugs such as AZD5991 (an Mcl-1 inhibitor), we observe very strong associations between drug response and gene effect in the drugs’ nominal targets. Correcting across all p-values risks amplifying these associations and overly correcting associations for non-targeted drugs.

Once the list of gene effects which are significantly predictive of drug response has been obtained, the number of dimensions which are linked to gene effects within the iCoVAE is selected by a simple heuristic: *k*_*s*_ = min (*n*_*sig*_, *L/*2), where *k*_*s*_ is the number of gene effects selected, *n*_*sig*_ is the number of significantly predictive gene effects (FDR *<* 0.05), and *L* is the dimension of the latent space. This ensures that features in the representations are only linked to significantly predictive gene effects, and that space is reserved in the latent space to encode information not related to gene effect, which is required for adequate reconstruction of gene expression profiles and prediction of drug response. In all cases where this method is employed, constraint selection is done using only the training set of the models (iCoVAE, PiCo), which utilise the auxiliary gene effect data, to avoid data leakage. This includes the holding out of cancer types in out-of-distribution generalisation experiments.

### 4.3 Cell line drug response prediction

We study the prediction performance of PiCo and compare baselines in two out-of-distribution settings. In the first (Fig. 3), PiCo is fitted on cell lines of selected cancer types in DepMap and GDSC, and we use this model to predict drug response on cell lines of new cancer types unseen during training. In this particular scenario, we fit the model using cell lines of more common cancer types, and test on uncommon cancer types. We include cancer types with at least 40 samples in the training set of the first stage of modelling, reserving all other cancer types for the test set. The cancer types selected for the training and test sets and the number of samples for each cancer type can be seen in Fig. B2. In this setting, we study 40 drugs, covering a range of small-molecule inhibitors and non-targeted treatments. In the second out-of-distribution setting (Fig. B4), we include samples of all cancer types except breast in the training set, and assess the performance of this PiCo model in drug response prediction on the unseen breast cancer cell lines. For this experiment, we focus on drugs which target important pathways in breast cancer and drugs which are commonly used in the treatment of breast cancer.

### 4.4 Treatment response prediction in breast cancer patients

To examine the use of iCoVAE representations to predict treatment response in a clinical setting, we form representations of gene expression data for breast cancer patients across the TransNEO cohort and its associated external validation set ARTemis+PBCP [29], again using a MAP representation. We then consider models to predict the residual cancer burden score (RCB score). We extend models predicting the more informative outcome, RCB score, to generate predictions of pathological complete response (pCR, equivalent to RCB score of zero) (Fig. B5).

For these models, we make use of both GDSC and CTRP data to form a consensus of gene effect-drug response relationships. We first select constraints separately for each of the drugs 5-fluorouracil, epirubicin (doxorubicin in CTRP), cyclophosphamide, and paclitaxel, then combine the selected constraints and the associations between the gene effect and the drug for each dataset. We then select constraints based on genes which are most often significantly associated with drug response across the 4 drugs in both datasets, and the strength of their association with drug response. We select 12 constraints using this method, to form a total of 16 constraints, with the anti-HER2 related genes *ERBB2, ERBB3, PIK3CA*, and *EGFR* also included. The 12 constraints selected using this method are *MCL1, PSMC1, FANCF, RAD1, PPM1D, SMU1, HSPA14, GBF1, MDM4, KNTC1, LRWD1* and *CEP97*.

Then, using these generated representations as an input, we fit ElasticNet-regularised linear regression models to predict RCB score, minimising mean squared error in an internal validation split using 10 individual 5-fold cross-validations each with a different random seed, in a grid search across the following hyperparameter search space, derived from [29]: regularisation parameter *C* in range {10^*−*4^, 10^4^}, L1:L2 penalisation ratio in range {0.0, 1.0}. We then evaluate the performance of the best models, as selected by cross-validation and refit on the entire TransNEO dataset, on the ARTemis+PBCP dataset, using root mean squared error (RMSE) and Spearman correlation as metrics for RCB score prediction. To assess the quality of the predictions of these models in inferring pCR (RCB= 0), we calculate the area under the receiver operating characteristic (AUROC) as a metric for pCR classification, using RCB score predictions directly.

### 4.5 Survival prediction in breast cancer patients

To examine the use of iCoVAE representations to predict survival in patients, we form representations of gene expression data for breast cancer patients across the SCAN-B [30] and TCGA [31]. In fitting the iCoVAE for this dataset, patient samples from SCAN-B for which there are no labels are used as additional unsupervised data in pretraining. TCGA samples are all unseen.

We then fit ElasticNet-regularised logistic regression models to predict 5-year breast cancer-free survival (BCFS) in these patients. We use the same cross-validation procedure and the same hyperparameter search space for the logistic regression, again aiming to minimise binary cross-entropy in a validation set.

In this setting, we use TCGA breast cancer data as an external validation set, and AUROC as a primary metric.

We also use a Cox proportional hazards model with *l*_1_ regularisation to predict BCFS. This model includes entire representations as well as clinical variables as features, compared to a model using only clinical variables and only representations. The same hyperparameter search space and cross-validation procedure as for the previously described regression models is used, minimising the negative log-likelihood. The concordance index (c-index) in the validation set of each fold of the cross-validation is then calculated as a performance metric, as well as the c-index for the same model evaluated on the TCGA external validation set.

In all models for SCAN-B and TCGA, clinical covariates are harmonised across the datasets. In particular, the clinical covariate for size is based on the American Joint Committee on Cancer Staging Manual for TNM (primary tumour, lymph nodes, metastases) staging of breast cancer. For TCGA, TNM staging information is available, but exact tumour sizes in millimetres are not; thus, we create a binary covariate valued 0 for T stages of Tis, T1a-c and 1 for T2-4, which therefore defines whether the primary tumour is greater than 20mm in size.

### 4.6 Permutation feature importance

To assess feature importance in PiCo prediction heads, we use a model-agnostic permutation-based feature importance method, as defined in [53]. In our implementation, to determine the importance for a feature, we randomly shuffle the feature 100 times and calculate the mean difference in performance between the model using the original input and the model using an input with the perturbed features. We repeat this for each random seed, and report the mean difference as the feature importance for the model using the given random seed. We either report a percentage change in correlation compared to the model using the non-perturbed input, Δ%*ρ*_*s*_, or a simple difference in a metric, e.g ΔAUROC.

### 4.7 Implementation

Hyperparameters (batch size, model size, learning rate, latent space dimension, dropout rate) for both the iCoVAE and VAE are selected through a 5-fold cross-validation procedure. Hyperparameter search for these models is implemented using the Tree-structured Parzen Estimator (TPE) implemented in the Optuna package, with 100 trials performed for each model [54], using a single random seed. These models are then refitted using 10 different random seeds to generate the pretrained encoders used in PiCo. Further details on the search space and model specifications can be found in Supplementary Information. Hyperparameters for each drug response prediction head are also optimised through a modified 5-fold cross-validation procedure—in this case, we run the hyperparameter optimisation separately with each of the 10 fitted iCoVAE/VAE models as a feature extractor, then refit these models on the combined training and validation data, to then evaluate their performance on the test data. All samples with both gene effect and drug response data available (i.e. present in both DepMap and GDSC) are always in the training set for both modelling stages to avoid data leakage, unless these samples are of held-out cancer types, in which case they are not present in any stage of training. The remaining samples, which have only drug response data available (in GDSC but not in DepMap), are split, with 80% added to the training data and 20% reserved for validation. Validation set performance is then reported by calculating the mean performance across the 5 folds and 10 random seeds. Where specified, performance is reported on a test set of samples held out across the entire training process. In this case, models with hyperparameters selected in the previously described steps are first fitted on the whole training and validation set for each of the 10 different random seeds and their associated best hyperparameters, with performance on the test set across the 10 final models calculated.

RNA-Seq gene expression data in units of log_2_(TPM + 1) and gene effect estimates from the Chronos model were downloaded from the DepMap portal^1^. For SCAN-B, TCGA, and TransNEO, gene expression values were converted from FPKM, TPM and raw counts, respectively, into the log_2_(TPM + 1) format.

In the clinical data studies, we filter the cell line gene expression data such that features used in the first stage of modelling are available for the clinical datasets, both of which provide RNA-Seq gene expression data. After this filtering step, we also remove genes for which expression has a variance of less than 1.0 (in log_2_(TPM + 1). The expression of these genes then makes up the input features for the iCoVAE and all subsequent models. Filtering gene expression by data availability and variance results in slightly different input features used for each experiment performed. For the experiments shown here, the input dimensions are: Depmap-GDSC: *d* = 6129, DepMap-TransNEO: *d* = 6519, DepMap-SCAN-B: *d* = 5137.

For clinical studies, the PiCo training process is carried out with no held-out cell line samples, such that the final models have been fitted on all cell lines with data available. These models are then fixed and used only to generate representations for patient data. New prediction heads are fitted using the generated representations for patients as inputs.

It was found in practice that the addition of further unlabelled cell line data to the training set for iCoVAE (i.e. moving from a supervised to a semi-supervised setting) slightly decreased performance in gene effect prediction on the supervised samples, although since the primary aim was improving representations of gene expression for downstream tasks, the decision was made to stay within a semi-supervised regime to maximise the quality of learned representations of gene expression.

Constraint selection is performed using the *statsmodels* Python package. All deep learning models are fitted in *PyTorch* version 2.0.1. All SVM, ElasticNet, and logistic regression models are fitted using the *scikit-learn* Python package. Cox models are fitted using the *lifelines* Python package [55].

## Acknowledgments

This work has been partly supported by UKRI grants MC UU 00002/16 and MC UU 00040/5 and the National Institute for Health Research (Cambridge Biomedical Research Centre at the Cambridge University Hospitals NHS Foundation Trust). O.M.R is also supported by a CRUK Career Establishment Award (RCCCEA-May22/100002). R.M. is supported by a CRUK Pre-doctoral Research Bursary (RCCPDB-May23/100003) and an NIHR Academic Clinical Fellowship. S-J.S. is a Lister Institute Prize Fellow and thanks Breast Cancer Now for funding his work as part of Programme Funding to the Breast Cancer

Now Toby Robins Research Centre. For the purpose of open access, the authors have applied a Creative Commons Attribution (CC BY) licence to any Author Accepted Manuscript version arising.

## 4.9 Data availability

I used data from DepMap version 23Q2, which is publicly available for download at https://depmap.org. I harmonised this with data from GDSC (October 2023 release), which is also freely available for download at https://cancerrxgene.org. The TransNEO and ARTemis+PBCP data used here can be downloaded from a GitHub repository provided in [29]. SCAN-B data is available in the dataset linked to [30]. TCGA data is freely available to download from https://cbioportal.org (Cell 2015 version).

## 4.10 Code availability

Code used to generate all results presented in this paper is available at https://github.com/domkirkham/pico.

## Appendix A Supplementary Information

### A.1 iCoVAE model

#### A.1.1 Model implementation and hyperparameter search space

The hyperparameter search space for the iCoVAE and VAE models is as follows:

- Batch size: {8, 16, 32, 64}
- Learning rate: {0.0001, 0.001, 0.01}
- Optimizer weight decay: {0.0001, 0.001, 0.01}
- Model size: {[512], [512, 256], [512, 256, 128]}
- Latent space dimension: {16, 32, 64}
- Dropout rate: {0.0, 0.1, 0.2, 0.3, 0.4, 0.5}

Model size provides the hidden layer widths for the encoder and decoder of the model. Adam [56] optimizer was used for all experiments. All deep learning models used LayerNorm, a ReLU activation and Dropout (where applicable).

#### A.1.2. Variance estimation for decoder, regressor, and conditional prior

In the model that the iCoVAE is based on, CCVAE [10], the constraints **s** are binary data and the distribution *q*(*s*^(*i*)^|*z*^(*i*)^) modelled as a Bernoulli distribution. Thus, a variance for this distribution does not need to be learned. For the conditional prior in CCVAE, which is assumed to be normally distributed, a single variance is learned for all dimensions in *z*, chosen for stability. To form an analogue for regression, where **s** is continuous, we need to define the variances of the regressor and the conditional prior carefully. We experimented with a range of richness in uncertainty estimation, with the richest representation involving a per-sample per-constraint variance, which was learned as a function of **s** or **z**, respectively. The second option required learning a per-constraint variance, which was shared by all samples. The first option resulted in model instability. Models trained with the per-target and shared variances were stable, but we observed that for the per-target variances, the model likelihood and performance focused on predicting constraints which were easier to predict from gene expression, ignoring other constraints which were more difficult to predict. This behaviour has been observed in calibrated decoders in VAEs.

Following this investigation we chose to model the variance of the observation distribution *p*_*θ*_(**x**|**z**) and the distribution *q*(**s**|**z**) as a single shared variance, which can improve quality of generations and naturally balances the reconstruction and KL-divergence terms in the VAE objective, as in the *β*-VAE [50].

##### A.2 PiCo improvements vary by cancer type

For oxaliplatin, PiCo provides the greatest benefit for cancers with infrequent *TP53* mutation. This effect is strongest in cell lines derived from cancers of the kidney, which are mostly composed of clear cell renal cell carcinomas (ccRCC) (PiCo-ElasticNet: RMSE = 1.486, *ρ* = 0.677, VAE-ElasticNet: RMSE = 1.580, *ρ* = 0.521, NN: RMSE = 1.629, *ρ* = 0.544, Fig. B1). In ccRCC cell lines, *TP53* is mutated in only 23.5% of samples, as opposed to 63.6% in pan-cancer [57, 58]. Dependency on *MDM4* should therefore be a useful predictor of resistance to platinum-based chemotherapy, since Mdm4-mediated inhibition of p53 is one of the primary mechanisms of p53 inactivation in ccRCC in the absence of inactivating *TP53* mutations.

PiCo also provides a large improvement in the prediction of AZD5991 response in liver cancer cell lines (PiCo-SVR: RMSE = 1.754, VAE-SVR: RMSE = 2.180, NN: RMSE = 2.065, Fig. B1c)—its nominal target Mcl-1 is highly expressed in the majority of hepatocellular carcinomas (HCC), and is a known survival factor in HCC. *MCL1* knockdown and/or inhibition has been observed to trigger apoptosis in in hepatocellular carcinoma [59]. As such, HCC is an active candidate for treatment with Mcl-1 inhibitors [60]. Although only providing a small benefit to overall drug response prediction for AZD5991 in the test set (PiCo-ElasticNet: *ρ* = 0.591, VAE-ElasticNet: *ρ* = 0.575, NN: *ρ* = 0.583), PiCo bases its predictions on features linked to *MCL1* and *MARCHF5* gene effect (*FI* = *−*17.68% and *FI* = *−*2.426%,Fig. B1b). A potential functional link between *MCL1* and *MARCHF5* has previously been highlighted [52].

##### A.3 Out-of-distribution generalisation to breast cancer cell lines

To further investigate out-of-distribution generalisation of PiCo in a more strict context, we instead held out all breast cancer cell lines, a more common but highly distinct cancer type (i.e. more strongly out-of-distribution), using data from cell lines of all other cancer types for model training. (Fig. B4). We see an improvement in prediction of ipatasertib response (PiCo-ElasticNet: *ρ* = 0.325, VAE-ElasticNet: *ρ* = 0.282, Fig. B4) and palbociclib response (PiCo-ElasticNet: *ρ* = 0.416, VAE-ElasticNet: *ρ* = 0.346), but a reduction in performance for the drugs AZD6378 (PiCo-ElasticNet: *ρ* = 0.081, VAE-ElasticNet: *ρ* = 0.363) and lapatinib (PiCo-ElasticNet: *ρ* = *−*0.070, VAE-ElasticNet: *ρ* = 0.241). Upon further investigation, we see that the iCoVAE fails to accurately infer the dependencies required for the prediction of response to these drugs in the unseen breast cancer cell lines, perhaps suggesting a reliance on spurious correlations to predict constraints in the training data, and further demonstrating the impact of constraint selection in PiCo. In the unseen breast cancer cell lines, prediction of response to palbociclib is reliant on latent dimensions encoding dependency on *CCND3, SKP2* and *N6AMT1*, which are all involved in the G0/G1 and S phases of the cell cycle [61, 62]. Palbociclib is an inhibitor of cyclin-dependent kinases *CDK4* and *CDK6*, which are crucial for the G1 to S transition. We emphasise again that in these experiments, the trained PiCo model has never seen data from breast cancer cell lines.

### Appendix B Supplementary Figures

**Fig. B1.**
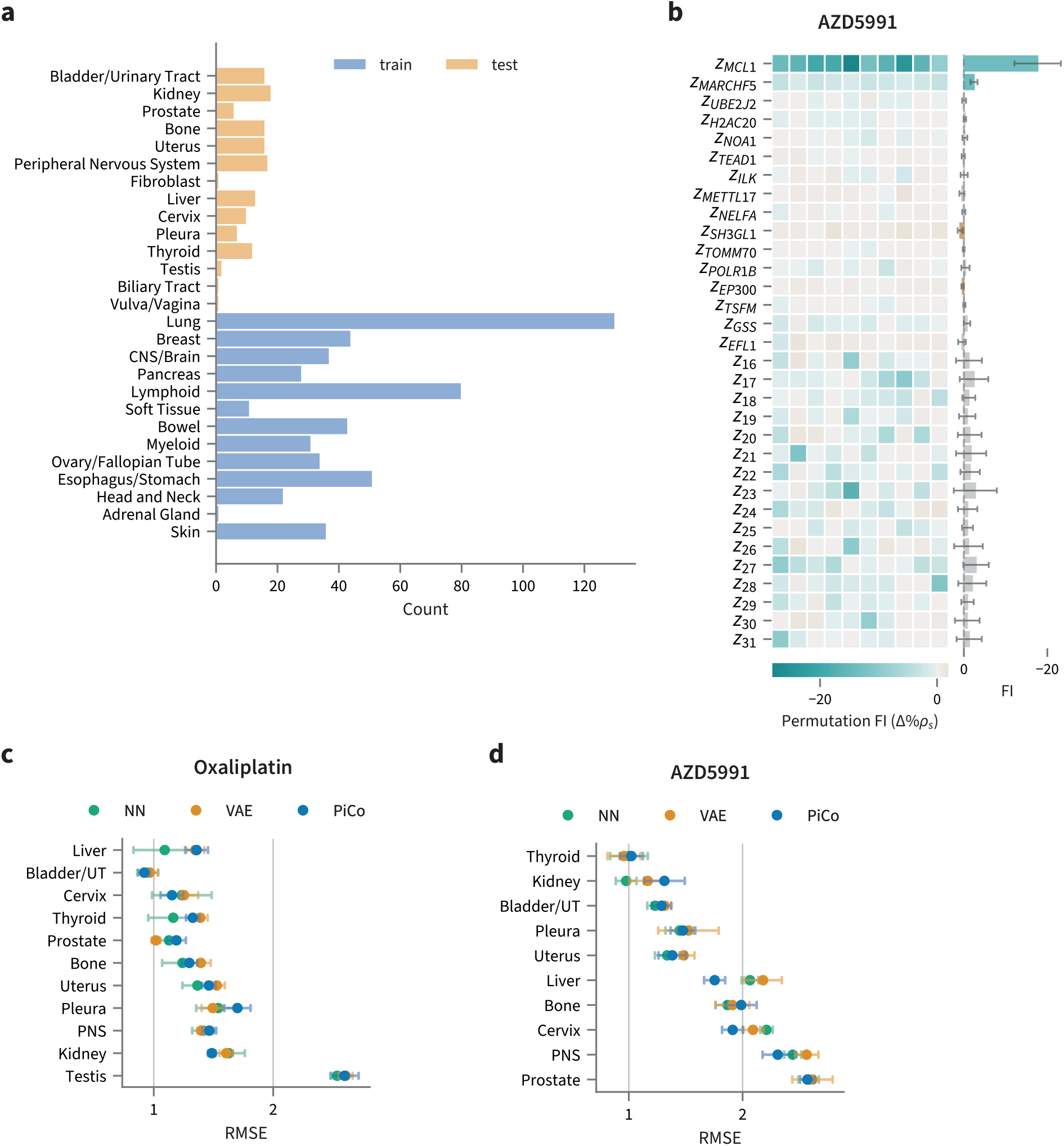
Additional information for Figure 3. **a** Number of samples of each cancer type in training and test sets for drug response prediction for the out-of-distribution prediction experiment with less common cancer types held out (16 cancer types with less than 40 samples in DepMap held out—only a subset of these samples are present in GDSC, such that some cancer types in the training set for this stage have fewer than 40 samples). **b** (left) Permutation feature importance (FI) for the model predicting AZD5991 response in test samples for each feature in PiCo-ElasticNet model, for 10 random seeds. (right) Marginal plot of FI across 10 random seeds. Error bars show +/- 1 s.d. Bars for features where perturbation consistently decreases performance are coloured blue. Bars are coloured grey if the mean FI +/- 1 s.d. includes zero FI. **c** Prediction performance in unseen cancer types for oxaliplatin, separated by cancer type. Spearman correlation, mean and +/- 1 s.d. across 10 random seeds shown by point and error bars. PNS: Peripheral nervous system. UT: Urinary tract. **d** as in **c** for AZD5991

**Fig. B2.**
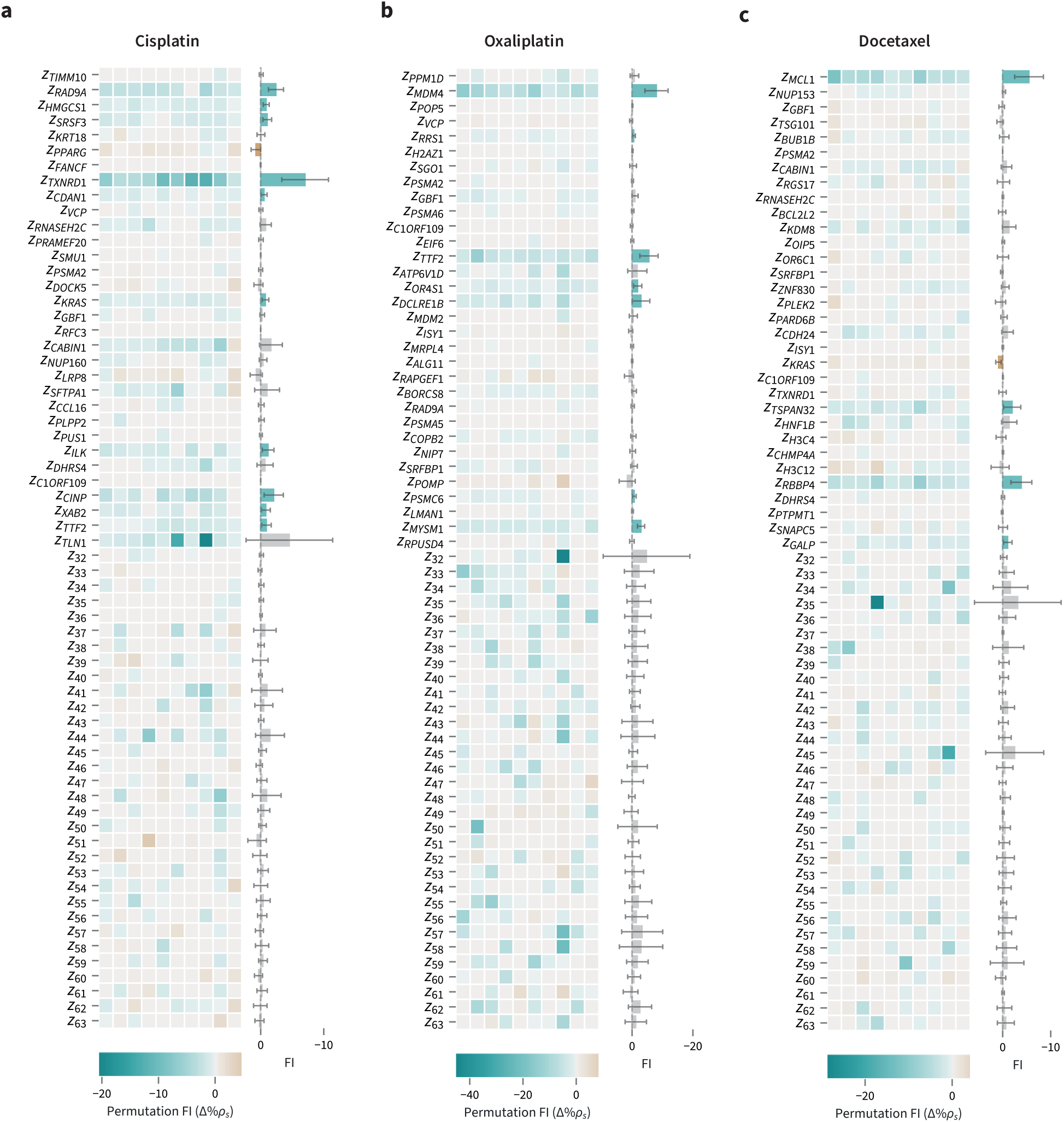
Additional permutation feature importance plots for PiCo on DepMap-GDSC. **a** (left) Permutation feature importance (FI) for the model predicting cisplatin response in test samples for each feature in PiCo-ElasticNet model, for 10 random seeds. (right) Marginal plot of FI across 10 random seeds. Error bars show +/- 1 s.d. Bars for features where perturbation consistently decreases performance are coloured blue. Bars are coloured grey if the mean FI +/- 1 s.d. includes zero FI. **b** as in **a**, for oxaliplatin. **c** as in a, for docetaxel.

**Fig. B3.**
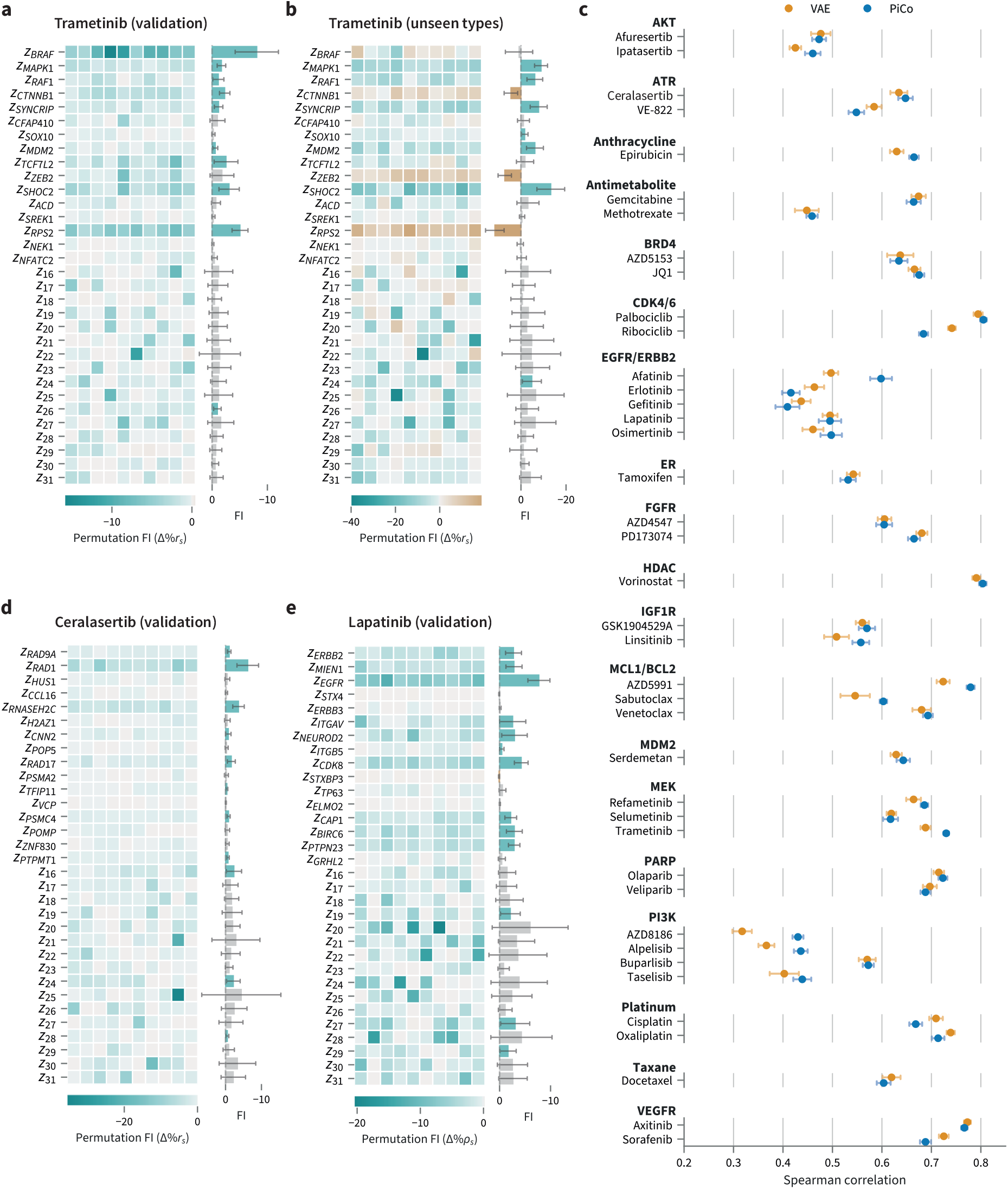
PiCo cross-validation performance on DepMap-GDSC. **a** Permutation feature importance (FI) for the model predicting trametinib response in validation samples for each feature in PiCo-SVR model, for 10 random seeds. (right) Marginal plot of FI across 10 random seeds. Error bars show +/- 1 s.d. Bars for features where perturbation consistently decreases performance are coloured blue. Bars are coloured grey if the mean FI +/- 1 s.d. includes zero FI. **b** as in **a** for the test set consisting of cancer types unseen during training. **c** Prediction performance in cross-validation, for a range of drugs. Drugs are grouped by nominal target, or targeted pathway. Mean Spearman correlation and +/- 1 s.d. across 10 random seeds. **d** as in **a** for ceralasertib. **e** as in **a** for lapatinib

**Fig. B4.**
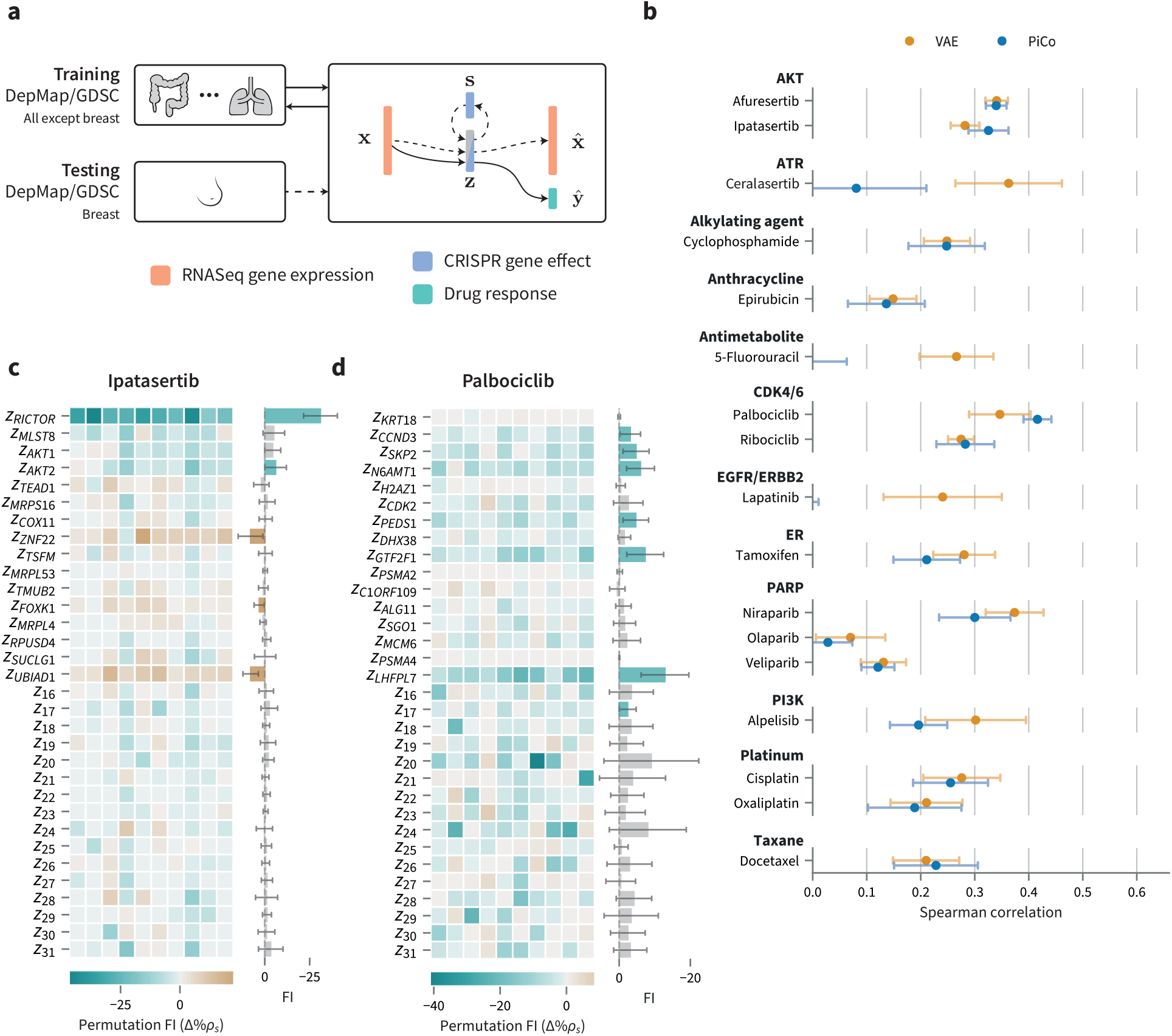
Robustness and feature importance in PiCo, with breast cancer cell lines held out. **a** To assess model generalisation, feature extractors and prediction models were trained on cell lines of all cancer types but breast, with samples of breast cancer type (*n* = 32) reserved for testing in zero-shot drug response prediction. **b** Prediction performance in unseen breast cancer cell lines, for a range of drugs. Drugs are grouped by nominal target, or targeted pathway. Mean Spearman correlation and +/- 1 s.d. across 10 random seeds. **c** (top) (left) Permutation feature importance (FI) for model predicting ipatasertib response in test samples for each feature in PiCo-ElasticNet model, for 10 random seeds. (right) Marginal plot of FI across 10 random seeds. Error bars show +/- 1 s.d. Bars for features where perturbation consistently decreases performance are coloured blue. Bars are coloured grey if the mean FI +/- 1 s.d. includes zero FI. **d** as in **c**, for palbociclib

**Fig. B5.**
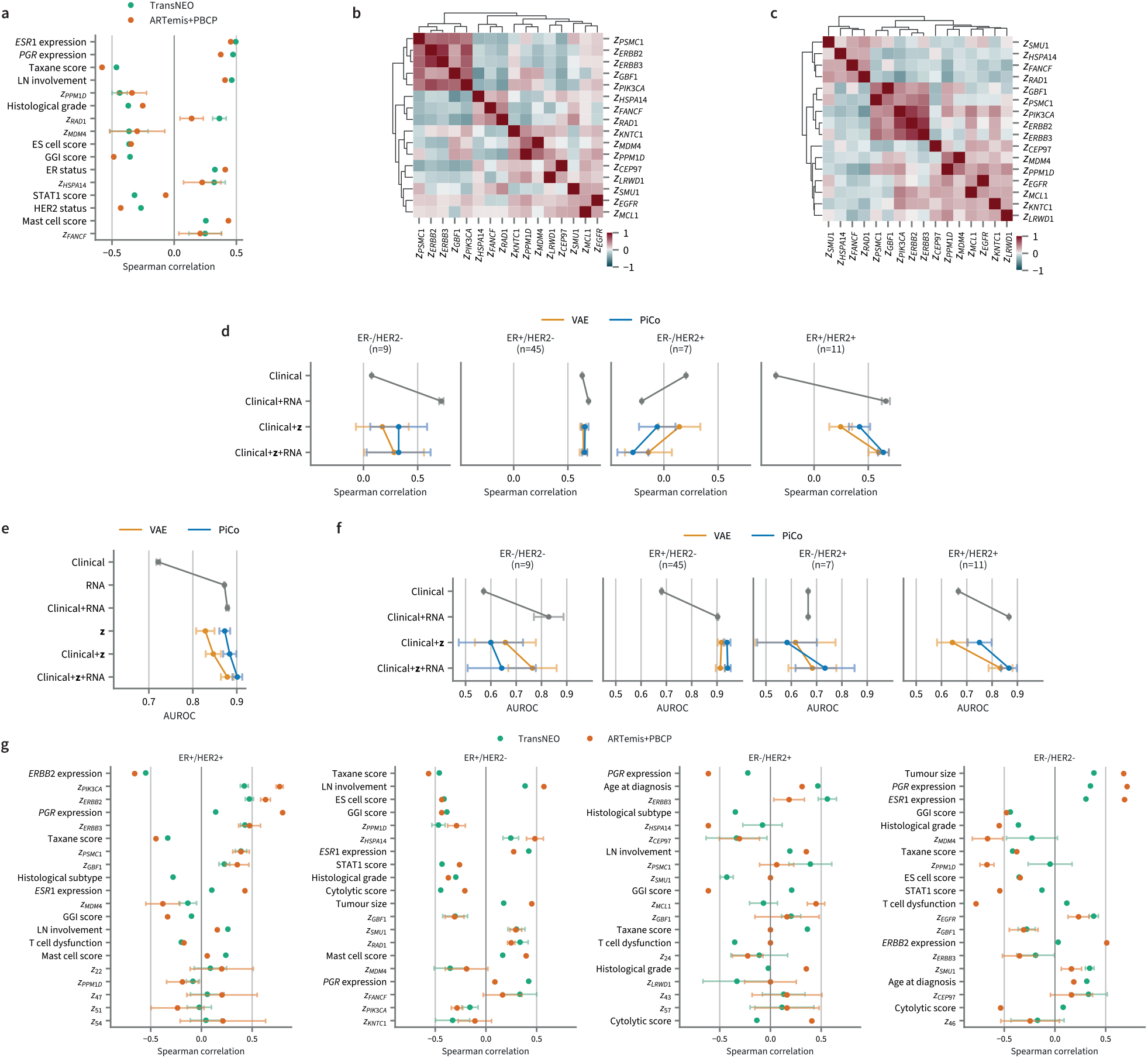
Additional information for TransNEO clinical study. **a** Single feature Spearman correlation of iCoVAE features with residual cancer burden (RCB) score in TransNEO **b** Correlation heatmap of iCoVAE representation features in TransNEO. **c** as in **b**, for the external validation set ARTemis+PBCP. **d** RCB score prediction performance in the external validation set ARTemis+PBCP, separated by molecular subtype. **e** Pathological complete response (pCR) prediction performance in the external validation set ARTemis+PBCP, using RCB score predictions directly. **f** as in **e**, separated by molecular subtype. **g** as in **a**, separated by molecular subtype. All performance and correlation are calculated over 10 random seeds. Mean +/- 1 s.d. shown by markers and error bars.

**Fig. B6.**
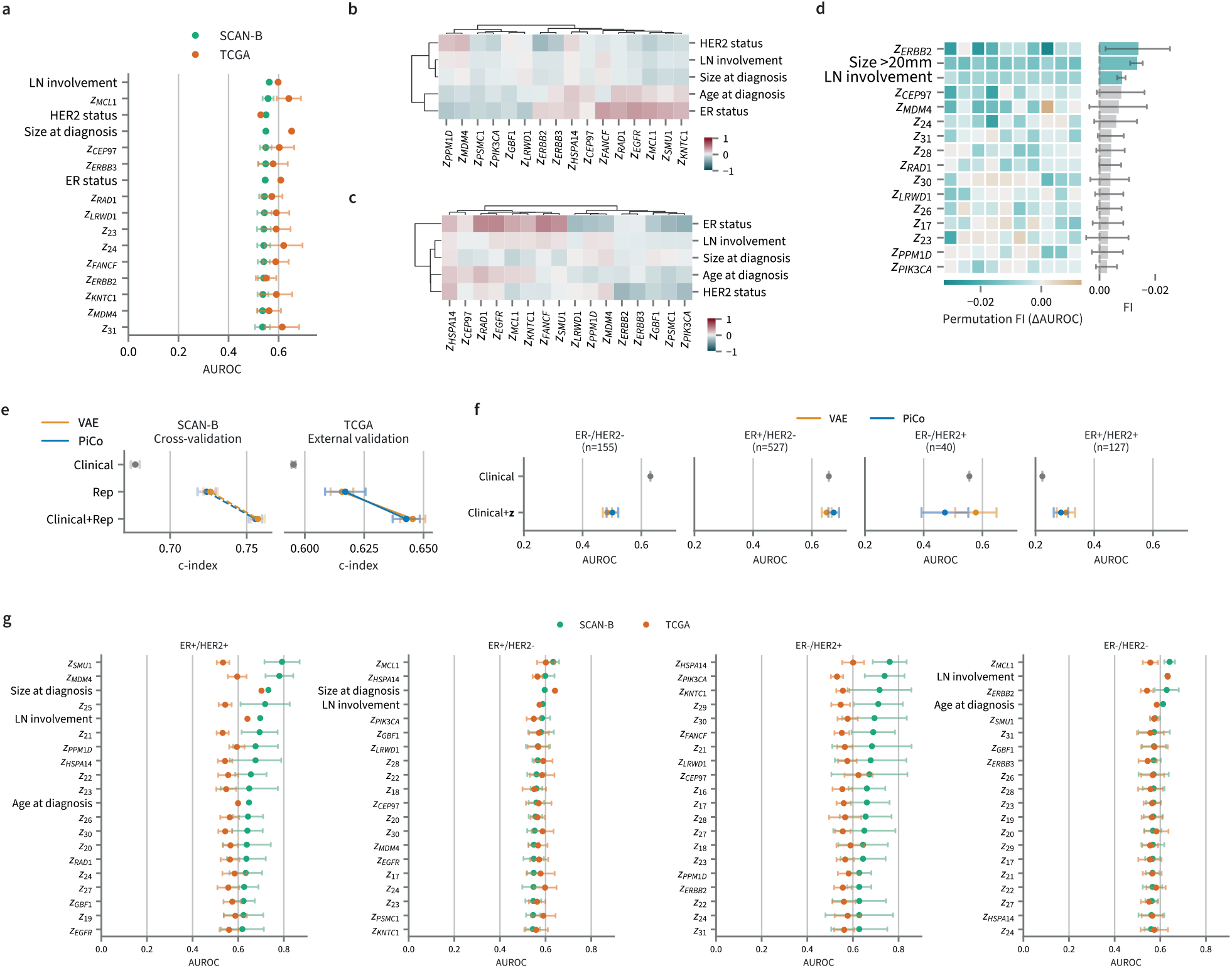
Additional information for SCAN-B clinical study. **a** Single feature AUROC of iCoVAE features with 5-year breast cancer free survival (BCFS) in SCAN-B and TCGA. **b** Correlation heatmap of iCoVAE representation features with clinical features available in SCAN-B. **c** as in **b**, for the TCGA external validation set. **d** Permutation feature importance for 5-year BCFS prediction performance on TCGA. **e** BCFS prediction performance using Cox model. **f** 5-year BCFS prediction performance in the TCGA external validation set, separated by molecular subtype. **g** as in **a**, separated by molecular subtype. All correlations are calculated over 10 random seeds. Mean +/- 1 s.d. shown by markers and error bars.

**Fig. B7.**
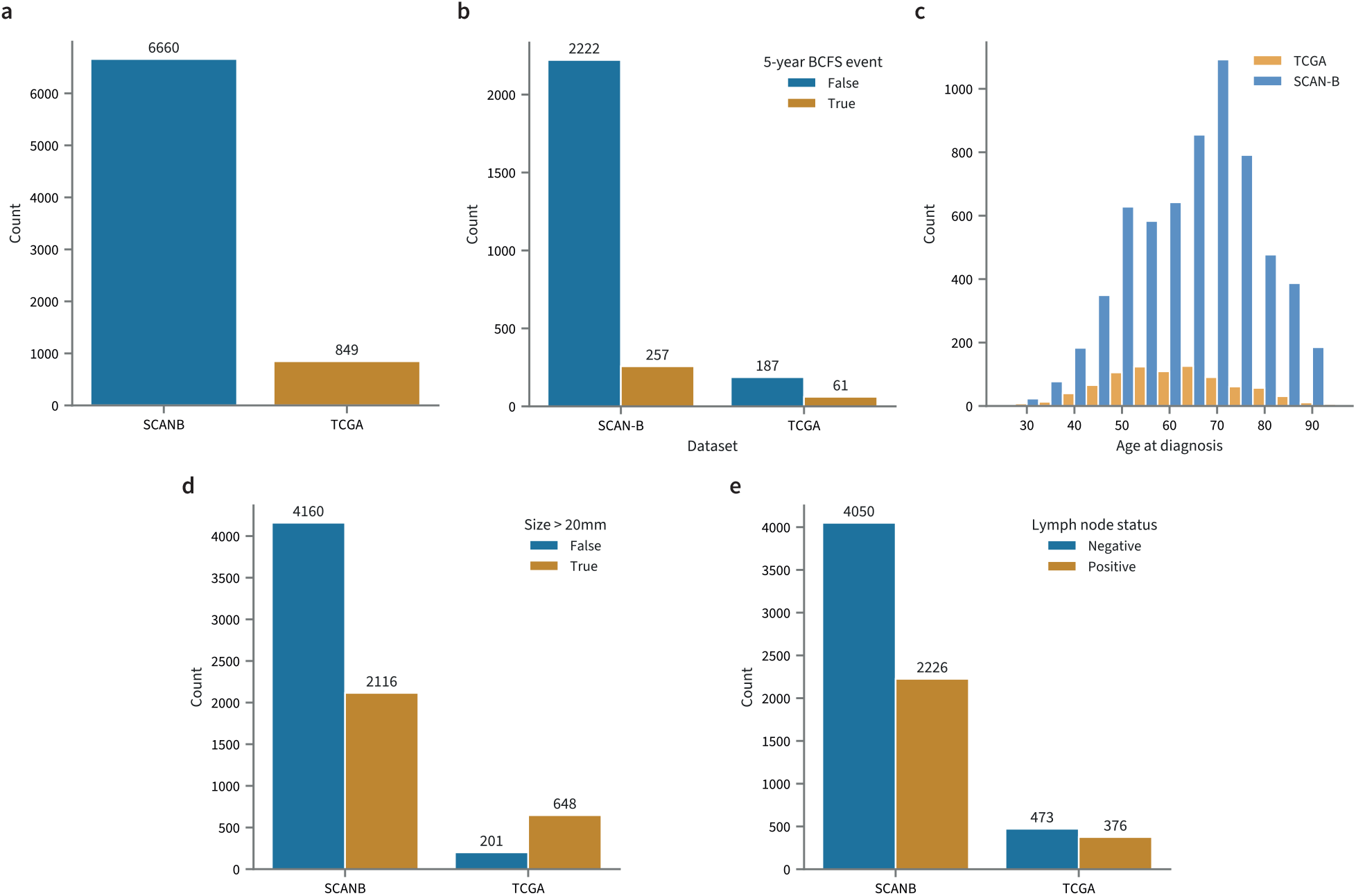
Comparison of SCAN-B and TCGA datasets. **a** Number of samples. **b** Number of BCFS events. **c** Age distribution of patients. **d** Number of tumours with size *>* 20mm. **e** Number of samples with positive lymph node status

## Footnotes

1 https://depmap.org

